# A model-guided pipeline for drug cardiotoxicity screening with human stem-cell derived cardiomyocytes

**DOI:** 10.1101/2021.09.10.459625

**Authors:** Alexander P. Clark, Siyu Wei, Trine Krogh-Madsen, David J. Christini

## Abstract

New therapeutic compounds go through a preclinical drug cardiotoxicity screening process that is overly conservative and provides limited mechanistic insight, leading to the misclassification of potentially beneficial drugs as proarrhythmic. There is a need to develop a screening paradigm that maintains this high sensitivity, while ensuring non-cardiotoxic compounds pass this phase of the drug approval process. In this study, we develop an *in vitro-in silico* pipeline using human induced stem-cell derived cardiomyocytes (iPSC-CMs) to address this problem. The pipeline includes a model-guided optimization that produces a voltage-clamp (VC) protocol to determine drug block of seven cardiac ion channels. Such VC data, along with action potential (AP) recordings, were acquired from iPSC-CMs before and after treatment with a control solution or a low-, intermediate-, or high-risk drug. We identified significant AP prolongation (a proarrhythmia indicator) in two high-risk drugs and, from the VC data, determined strong ion channel blocks that led to the AP changes. The VC data also uncovered an undocumented funny current (I_f_) block by quinine, which we confirmed with experiments using a HEK-293 expression line. We present a new approach to cardiotoxicity screening that simultaneously evaluates proarrhythmia risk (e.g. AP prolongation) and mechanism (e.g. channel block) from iPSC-CMs.

## INTRODUCTION

In the 1990s, cardiotoxicity was the number one cause for the US Food and Drug Administration to withdraw or restrict the use of a drug on the market (Lasser et al., 2002). Such drugs were identified because they increased the prevalence of lethal heart arrhythmias (Lasser et al., 2002; Roden, 2005). It was found that many of these drugs block the human ether-à-go-go related gene (hERG) channel, which is responsible for conducting repolarizing potassium current (I_Kr_) and is known to be protective against the development of arrhythmias.

These findings inspired the development of hERG-based screening approaches (EMA, 2005; Windley et al., 2018; Yang et al., 2020) that have essentially eliminated the risk of lethal proarrhythmic drugs making it to market (Sager et al., 2014). The high sensitivity of these approaches comes at the cost of low specificity (De Bruin et al., 2005; Gintant, 2011; Hancox et al., 2008), leading to false positive classification for some safe and effective therapies, like verapamil and ranolazine (Crumb et al., 2016; Johannesen et al., 2014). Such misclassified drugs counteract the proarrhythmic effects on hERG by blocking ion channels that conduct current in the opposing direction (e.g. depolarizing calcium and sodium channels), emphasizing the need for multi-channel screening.

To address the specificity shortcomings of hERG-based approaches, the Comprehensive *in Vitro* Proarrhythmia Assay (CiPA) initiative was started in 2013 to guide the development of more accurate preclinical tests (Sager et al., 2014). The group established a three-step drug screening pipeline that includes: 1) quantifying drug effects on multiple ionic currents using expression line cells, 2) integrating these effects into *in silico* models and using them to evaluate a drug’s proarrhythmic potential, and 3) validating simulations with human-derived induced pluripotent stem cell cardiomyocytes (iPSC-CMs) and human ECG studies.

The CiPA initiative identified 28 drugs with known clinical characteristics for testing and validating new screening methods. These drugs were categorized into low-, intermediate-, and high-risk groups based on their risk of causing lethal arrhythmias. In 2016, the dose-response behavior for these drugs was determined on seven cardiac ion currents (Crumb et al., 2016). By applying this multi-channel drug block data to *in silico* models, drug cardiotoxicity risk has been accurately predicted at the single-cell, tissue, and whole heart levels (Costabal et al., 2019; Gong and Sobie, 2018; Passini et al., 2017; Sahli-Costabal et al., 2020; Tomek et al., 2019; Zhou et al., 2020). These approaches were also used to evaluate the proarrhythmic potential of COVID-19 therapies, such as azithromycin and hydroxychloroquine (Sutanto and Heijman, 2020; Varshneya et al., 2021; Whittaker et al., 2021).

In parallel to these *in silico* studies, human-derived iPSC-CMs have become a standard *in vitro* tool to measure surrogate markers of proarrhythmic risk, such as AP prolongation and changes in calcium transients (Bedut et al., 2016; Charrez et al., 2021; Klimas et al., 2016; Kopljar et al., 2018; Mathur et al., 2015). While approaches that use these markers are efficient for predicting proarrhythmic risk, they do not provide insight into the mechanism of action of a drug. Recently, steps have been taken to address this shortcoming by fitting *in silico* models to iPSC-CM AP and calcium data to predict drug block of I_CaL_, I_Kr_, and I_Na_ (Jæger et al., 2021a, 2021b).

For all the value that iPSC-CMs have provided to the drug screening process over the last decade, they are still an imperfect model of adult physiology, with an immature phenotype, high degree of heterogeneity, and depolarized maximum diastolic potential, leading to spontaneous APs (Goversen et al., 2018b). These features make it difficult to record consistent and reliable measures of adult proarrhythmic risk. Dynamic clamp has been used to address some of these shortcomings by injecting a synthetic hyperpolarizing I_K1_-like current to stop the spontaneous beating and establish a stable maximum diastolic potential below −70mV (Fabbri et al., 2019; Quach et al., 2018). When paced from this hyperpolarized resting membrane potential, cells have a more consistent, and adult-like AP phenotype, making drug-induced AP changes easier to interpret (Goversen et al., 2018a; Li et al., 2019).

In this study, we aim to build on the approaches outlined above to develop a novel pipeline that determines both drug-induced cardiotoxicity risk and mechanism from iPSC-CMs. Specifically, we use automated experiment design to develop an iPSC-CM voltage clamp (VC) protocol. The optimized 9-second protocol was used to identify which of seven ionic currents (I_Kr_, I_CaL_, I_Na_, I_to_, I_K1_, I_f_, and I_Ks_) were strongly blocked by drugs selected from each of the three CiPA risk groups: low (verapamil), intermediate (cisapride), and high risk (quinidine, quinine). These drugs were selected to determine whether the protocol could identify ion channel block for single-channel (e.g. cisapride block of I_Kr_ at >15x EFPC) and multi-channel drugs (e.g. verapamil, quinine, and quinidine all at 3x EFPC). In contrast to previous approaches with iPSC-CMs, this short-duration protocol enabled us to investigate drug block of seven channels within a single cell. We show that these drug targets (e.g. quinine block of I_Kr_) can explain the AP morphology changes (e.g. AP prolongation) seen after treatment with a drug. With the VC protocol, we also identified a previously unreported block of funny current (I_f_) by quinine at 3x the effective free therapeutic plasma concentration (EFPC), which was confirmed with a dose-response study on a HEK-293 cell line stably expressing HCN1.

To the best of our knowledge, we have developed the first VC protocol optimization algorithm that is specifically designed for the purpose of identifying multi-channel drug block in an electrically excitable cell. We believe that the pipeline outlined in this paper has the potential to improve cardiotoxicity risk assessment and, ultimately, increase the number of safe and effective drugs available to patients.

## RESULTS

### An in silico-in vitro pipeline to determine cardiotoxicity risk and mechanism (Figure 1)

The first step in the pipeline is to use an *in silico* iPSC-CM model-guided genetic algorithm to design a voltage-clamp protocol that isolates individual currents (Step 1). While the voltage-clamp protocol could in principle be designed to isolate any of the ionic currents present in the *in silico* model, in this study we focused on seven currents that are most associated with AP morphology (I_Kr_, I_CaL_, I_Na_, I_to_, I_K1_, I_f_, I_Ks_). Optimized voltage-clamp, as well as spontaneous and I_K1_ dynamic clamp and paced AP data, is acquired from a patient-derived iPSC-CM before and after drug application (Step 2). The I_K1_ dynamic clamp data is used to measure surrogate markers of cardiotoxicity (Step 3), while the optimized voltage-clamp data is used to identify ion channel targets (Step 4). Dose-response data is then acquired for the identified targets using expression line cells (Step 5). For example, in this study we acquired the dose-response data for quinine block of HCN1, which further validated our findings on the unreported block of I_f_ by quinine.

**Figure 1:**
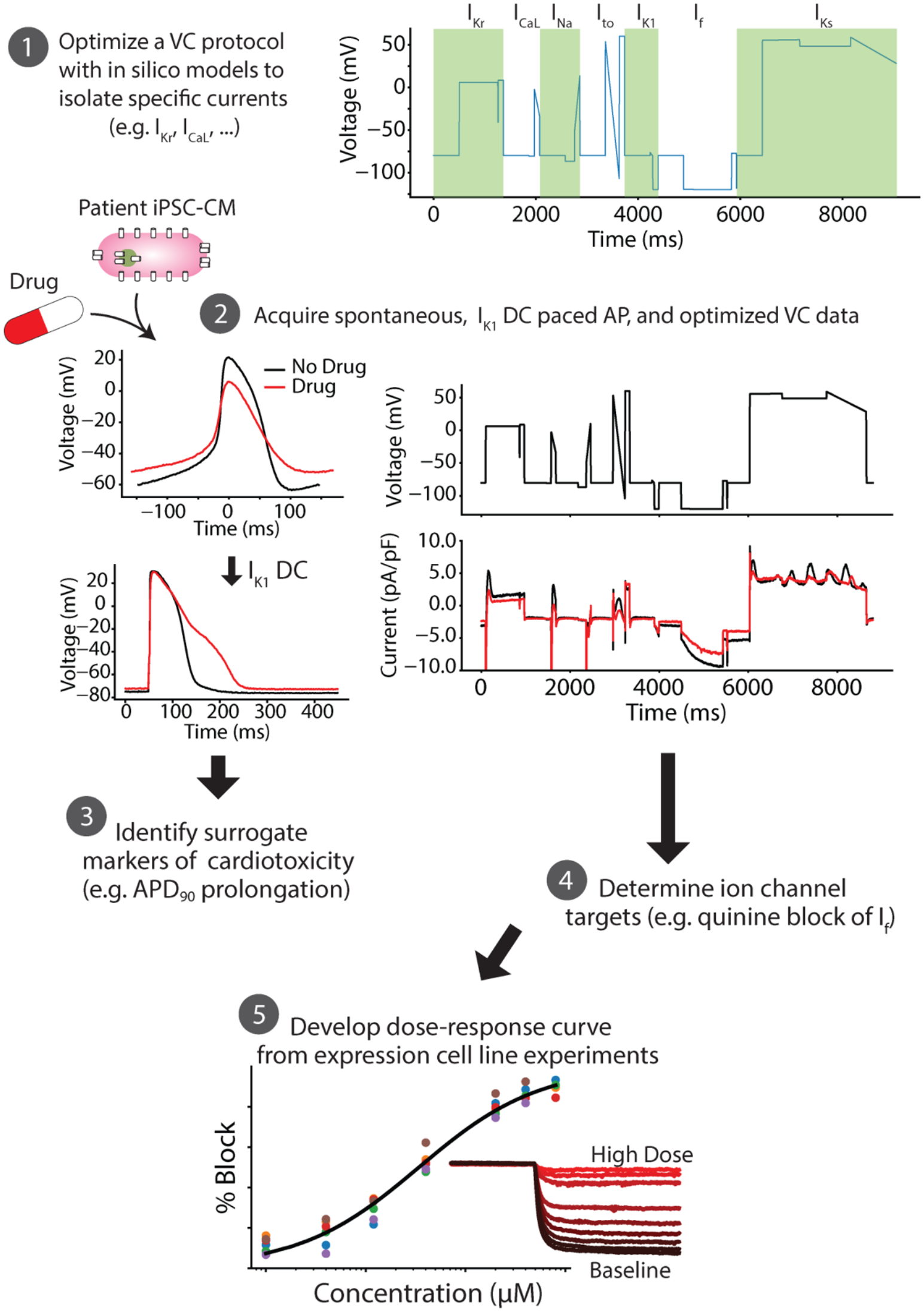
An *in silico-in-vitro* pipeline to determine drug cardiotoxicity risk and mechanism. Step 1, The Kernik-Clancy model with experimental artifacts is used to develop a voltage clamp protocol that is specifically designed to isolate currents. **Step 2,** Spontaneous, I_K1_ dynamic clamp and paced AP, and optimized voltage-clamp data is acquired from a patient-derived iPSC-CM before and after drug application. **Step 3,** The change in I_K1_ dynamic clamp and paced AP data from pre-to post-drug application is used to identify AP prolongation, a surrogate marker of cardiotoxicity. **Step 4,** Changes in voltage-clamp data is used to determine the ion channels targeted by the drug. **Step 5,** After identifying the ion channel targeted by the drug, a dose-response curve is developed for each of these ion channels using expression line cells.

### Optimizing a VC protocol to isolate individual currents for drug cardiotoxicity screening

We used a model-guided experimental design approach to optimize a VC protocol that isolates the contribution of individual currents at different timepoints. Our rationale for isolating current contributions, similar to our previous work (Groenendaal et al., 2015), is that this enables tracking changes in individual currents during iPSC-CM drug studies and using these changes to identify ion channel targets.

We used the recent Kernik-Clancy iPSC-CM model (Kernik et al., 2019) in the optimization algorithm to simulate a cell’s response to VC protocols. Because we expected the protocol to isolate currents during segments that are most sensitive (e.g., <10ms after a voltage step) to experimental artifacts (see Materials and Methods), we added equations that incorporate these effects (e.g. voltage offset, leak current, and series resistance), as was recently shown to be critical (Lei et al., 2020).

We used the genetic algorithm to optimize VC protocols that maximize the current contribution for one of seven ionic currents: I_Kr_, I_CaL_, I_Na_, I_to_, I_K1_, I_Ks_, and I_f_ (Figure 2A, Appendix – Figures 2-8). The durations of the seven protocols ranged from ∼1400 ms for I_to_ to ∼3800 ms for I_Ks_. The VC protocols for each current were systematically shortened (see Materials and Methods) and then combined into a single protocol with the segments that maximized the isolation of each ionic current (Figure 2B). The resulting optimized protocol was just over 9 seconds, i.e., short enough to be run multiple times in each cell both under control conditions and with drug application.

**Figure 2:**
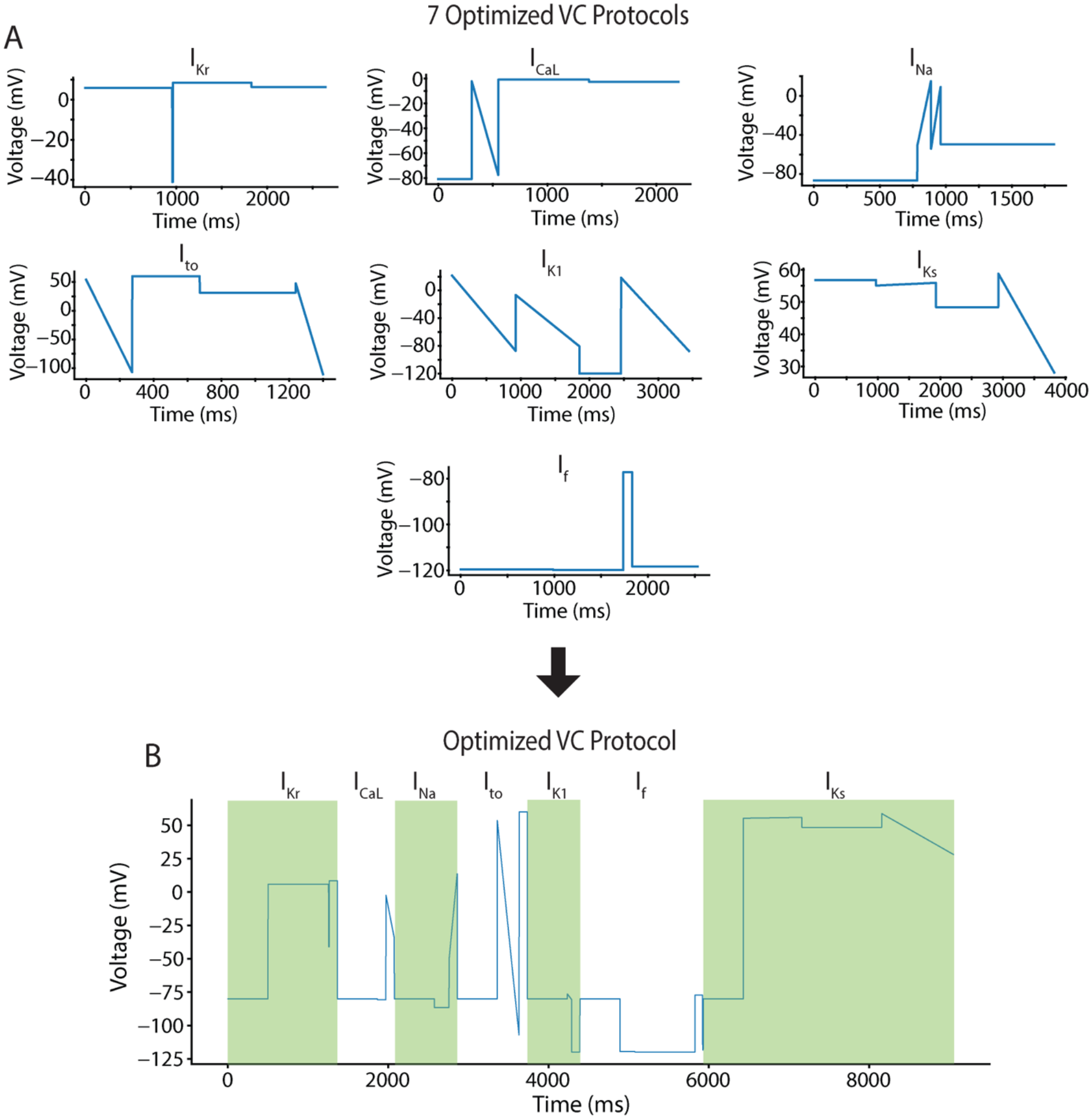
A genetic algorithm optimized a voltage clamp protocol to isolate individual current contributions in an *in silico* iPSC-CM model. **A,** The genetic algorithm was used to develop individual voltage clamp protocols for seven of the most predominant ionic currents in cardiac cells. **B,** These seven protocols were systematically shortened and joined together by a −80mV holding step to produce an optimized voltage clamp protocol that isolates each of the seven currents.

We validated the VC protocol by applying it to a different iPSC-CM model (Paci et al., 2018) and comparing the windows of maximum current. This step provided us with confidence that the VC protocol could isolate currents during the same time windows from a cell with different conductances and kinetics (Appendix – Figure 9).

### Synthetic maturation of iPSC-CMs by I_K1_ dynamic clamp improves interpretability of iPSC-CM AP data

We conducted *in vitro* experiments using isolated iPSC-CMs. The dynamically clamped and paced APs were acquired by injecting a synthetic I_K1_ model current (Ishihara et al., 2009) into the cells until spontaneous AP generation stopped and the resting membrane potential was below −65 mV.

The iPSC-CMs displayed a heterogeneous phenotype (Figure 3A), which is consistent with previous work on single-cell iPSC-CMs (Garg et al., 2019). Figure 3A shows the spontaneous behavior of six cells that were selected to highlight the heterogeneity in the population. By dynamically clamping I_K1_ and pacing, we were able to elicit APs from all these cells (Figure 3B). This finding is consistent with previous work showing the value of I_K1_ dynamic clamp in reducing (although not eliminating) cell-to-cell heterogeneity while eliciting more adult-like AP behavior from iPSC-CMs (Li et al., 2019). The I_K1_ dynamic clamp and paced APs (n=40) had an average resting membrane potential of −74.2 ± 2.8 mV and action potential duration at 90% repolarization of 142.0 ± 48.3 ms.

**Figure 3:**
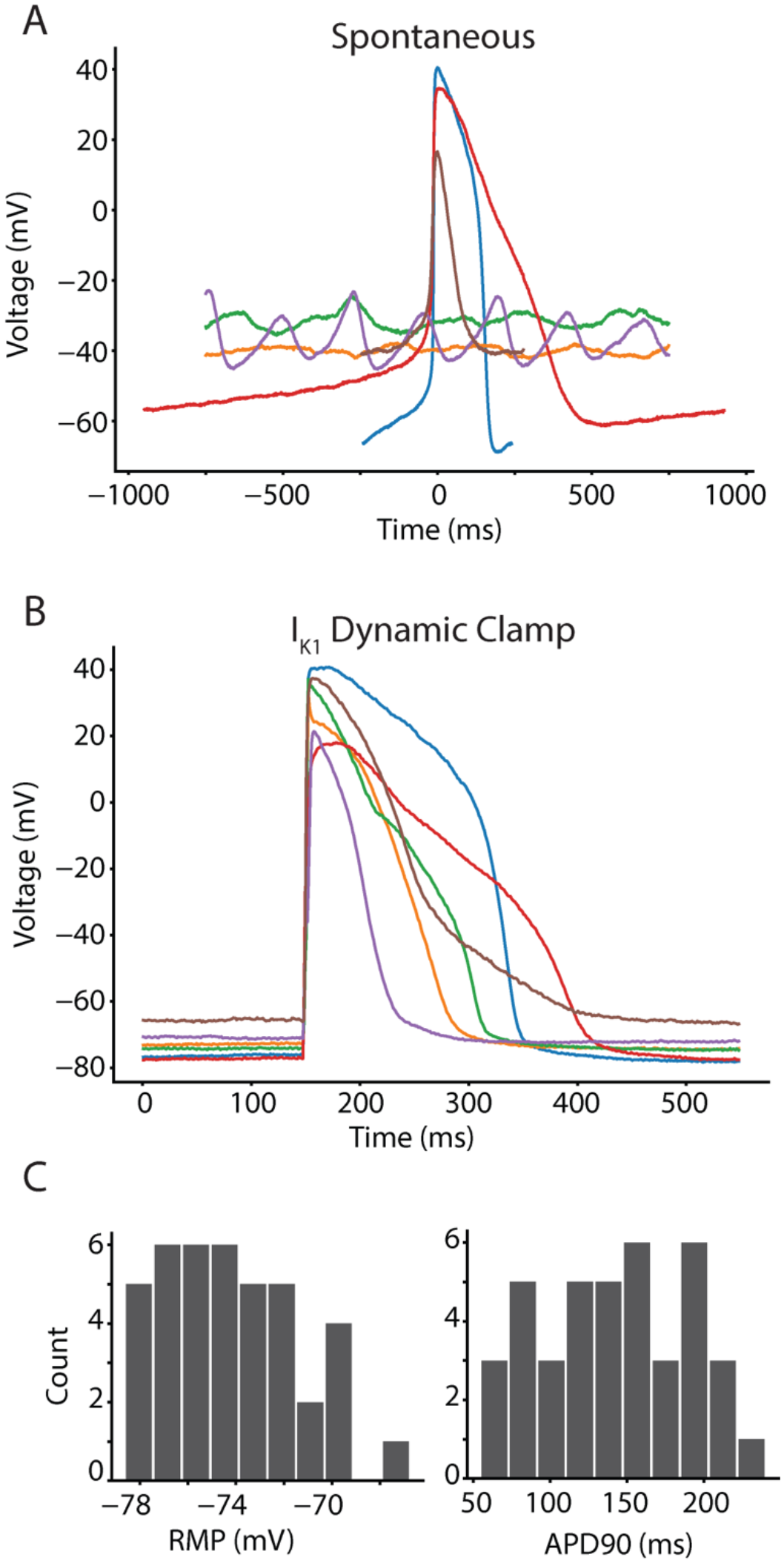
Synthetic maturation of iPSC-CMs with dynamically clamped I_K1_ ensures consistent APs under 1Hz pacing conditions. **A**, Spontaneous behavior from six cells indicates the level of intercell heterogeneity and inconsistent presence of APs in cell population. **B**, I_K1_ dynamic clamp applied to the same cells makes them appear more mature when paced at 1Hz. **C**, Histograms for the I_K1_ dynamically clamped and paced AP resting membrane potential (−74.2 ± 2.8 mV) and action potential duration at 90% repolarization (142.0 ± 48.3 ms) for the 40 cells in this study.

### I_K1_ dynamic clamp AP data identifies surrogate markers of cardiotoxicity

We compared changes in AP features for iPSC-CM data acquired before and after application of cisapride (n=6), verapamil (n=9), quinidine (n=6), quinine (n=9), or DMSO control (n=10). Table I shows the percent block of each cardiac ion channel by these drugs based on previous results (Crumb et al., 2016). Cisapride is a CiPA-labeled intermediate-risk drug and blocks I_Kr_ strongly and specifically at the concentrations used in this study. The cell in Figure 4A shows AP prolongation after cisapride treatment that is characteristic of such I_Kr_ block. Verapamil is a low-risk drug that moderately blocks I_CaL_ and lightly blocks I_Kr_ at the concentrations used in this study. Figure 4B shows a verapamil-treated cell that displays AP shortening after verapamil treatment. Most of the shortening in 4B appears to be due to AP triangulation, a morphological change also seen in experimental data from human cardiomyocytes (Britton et al., 2017). Quinidine and quinine are both high-risk CiPA drugs that block multiple ion channels with a strong affinity for I_Kr_. Both quinidine-(p=.034, Figure 4C) and quinine-treated (p=.0003, Figure 4D) cells show AP prolongation after drug application.

**Figure 4:**
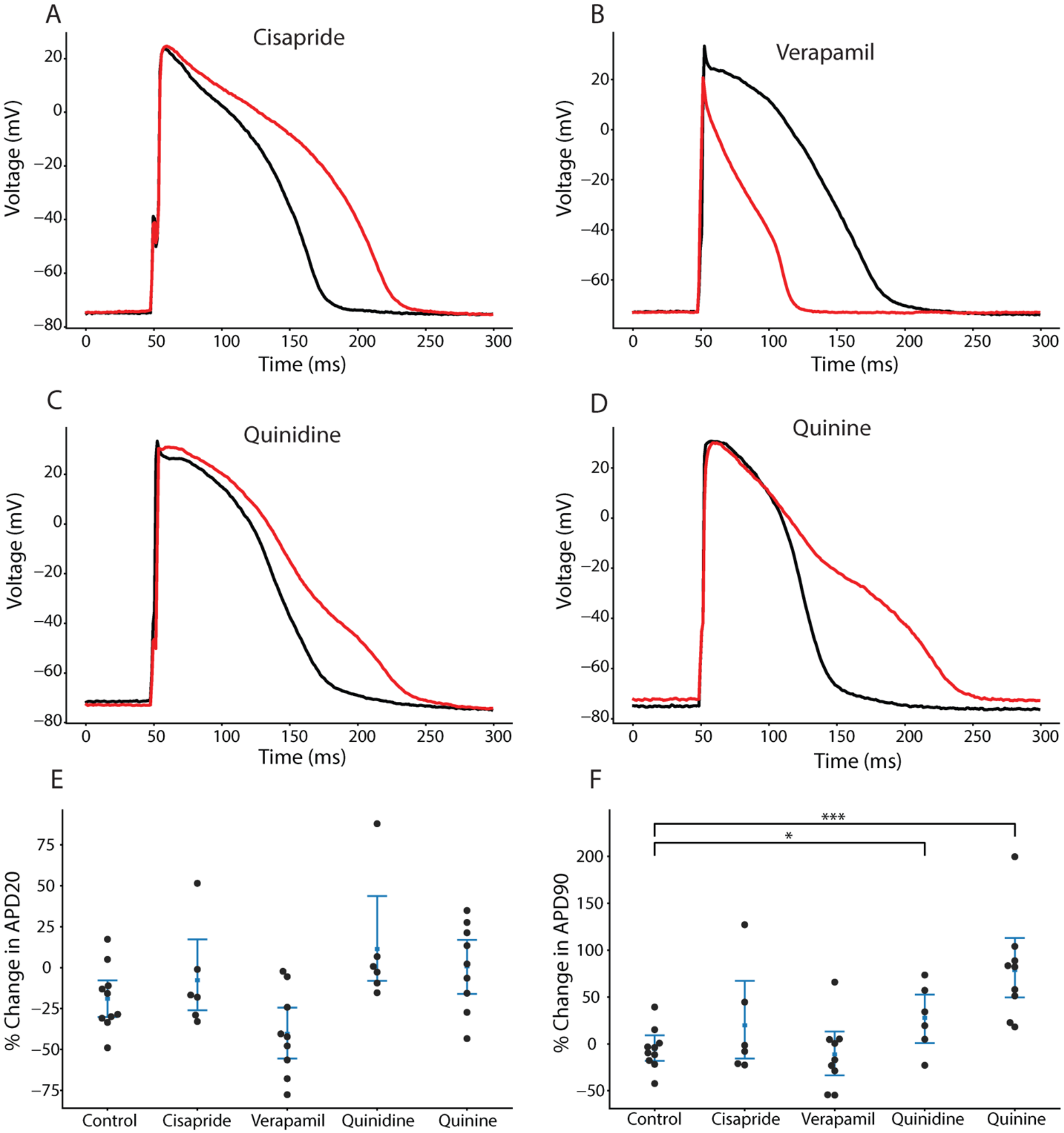
Drug treatment resulted in AP morphology changes consistent with the effects caused by drug block of the expected currents. Four different cells with dynamically clamped I_K1_ and paced at 1Hz under treatment with cisapride (**A**), verapamil (**B**), quinidine (**C**), and quinine (**D**). **E**, No cells showed a significant change in APD20 relative to DMSO control. **F**, Change in APD90 relative to DMSO control. Quinidine (p=.034) and quinine (p=.0003) showed significant prolongation.

**Table I:** The optimized VC protocol correctly identifies strong drug blocks

| Drug | $I_{Kr}$ | $I_{CaL}$ | $I_{Na}$ | $I_{to}$ | $I_{K1}$ | $I_f$ | $I_{Ks}$ |
| --- | --- | --- | --- | --- | --- | --- | --- |
| **Cisapride (125 nM)** | 95%* | 1% | 2% | 13% | 5% | ?? | 2% |
| **Verapamil (150 nM)** | 21% | 39% * | <1% | 1% | 3% | ?? | 3% |
| **Quinidine (2,529)** | 89%* | 16% | 10% | 43%* | 1% | ?? | 27% |
| **Quinine (12,000 nM)** | 72%* | 29% | 28% | 15% | <1% | 32%*† | 20% |

The percent block data in this table is taken from Crumb et al. (2016). The optimized VC protocol correctly identifies drug blocks predicted (from Crumb et al) to reduce currents by greater than 30% (\*) and identifies funny current as a novel target of quinine (†).

**Table II:** Kernik-Clancy and experimental artifact equations used in GA optimization.

|  |  |
| --- | --- |
| $I_{ion} = I_{Na} + I_{CaL} + I_{Kr} + I_{Ks} + I_{K1} + I_{to} + I_f + I_{CaT} + I_{NCX} + I_{PMCA} + I_{NaK} + I_{bCa} + I_{bNa}$ | Kernik-Clancy iPSC-CM Model |
| $I_{leak} = g_{leak}V_m$ | Seal leak current |
| $I_{out} = I_{ion} + I_{leak}$ | Observed current |
| $V_p = V_{cmd} + \alpha R_s I_{out}$ | $R_s$ compensation |
| $\frac{dV_m}{dt} = \frac{1}{R_s C_m} (V_p + V_{off} - V_m) - \frac{1}{C_m} I_{out}$ | Membrane voltage differential |

**Table III:**
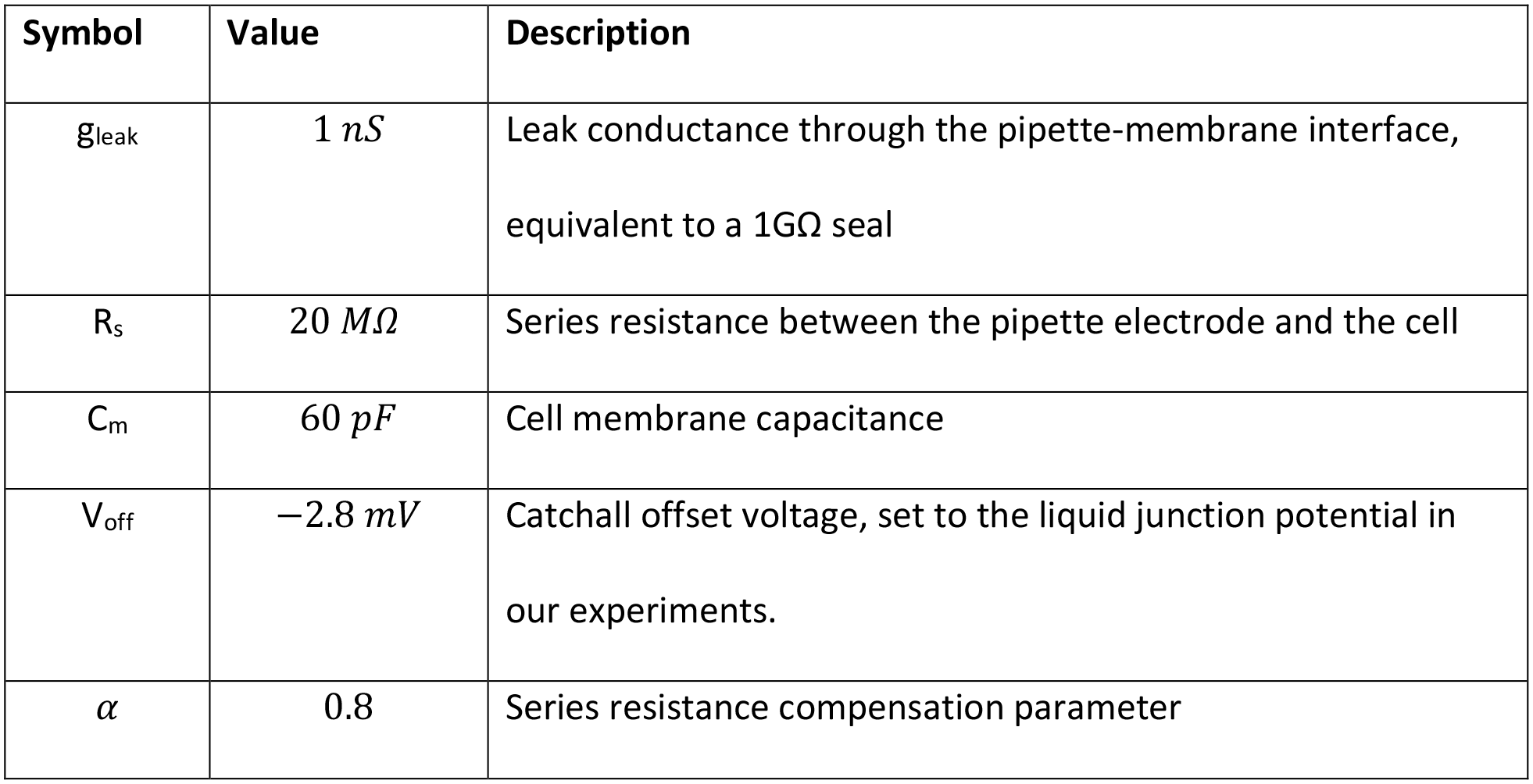
Experimental artifact parameters values used in GA simulation.

Verapamil-treated cells showed shortening in the action potential duration at 20% of repolarization (APD_20_), but the difference between these and control cells was not statistically significant (p=.056, Figure 4E). Cells that were treated with quinidine and quinine showed a significant prolongation in the action potential duration at 90% of repolarization (APD_90_) when compared to control cells (Figure 4F). These findings agree with the CiPA classification for verapamil (low risk), quinidine (high risk), and quinine (high risk). Because APD_90_ prolongation is an established proarrhythmia risk indicator, these data correctly suggest that quinidine and quinine have proarrhythmic potential at 3x their EFPC. Cisapride-treated cells did not show a significant change in APD_90_, despite it being a stronger I_Kr_ blocker than quinidine and quinine at the concentrations used in this study. This may be because quinidine and quinine also block a small amount of I_Ks_ at the concentrations used in this study. Because these drugs block both dominant repolarizing potassium currents (I_Kr_ and I_Ks_), they may appear to cause greater and more consistent AP prolongation in such a heterogeneous iPSC-CM population.

### The optimized VC protocol qualitatively identifies drugs that block greater than 30% of an ionic current

While the AP data described above was used to identify surrogate markers of drug cardiotoxicity, it did not provide specific information about the drugs’ mechanism. To identify the targets of each drug, we compared the average change in VC responses for a given drug to the average change of cells treated with DMSO control. Because the protocol was designed to isolate one specific ionic current during a brief window of the protocol, differences of the changes in each of these seven segments could reveal ionic mechanism.

As a demonstration of this approach, Figure 5A shows a representative example of a cell that was treated with quinine. The zoomed-in panel in Figure 5B shows the portion of the VC protocol that maximized the current contribution of I_Kr_ relative to the total current. This inset shows a decrease in the total current present after treatment with quinine throughout the time window. The difference is particularly pronounced from 860 ms to 865 ms and from 870 ms to 875 ms, which corresponds to the segments where I_Kr_ is predicted by the model to have a large relative contribution compared to the other ionic currents present. According to our modeling work, the 865-870 ms segment is predicted to have the largest amount of I_Kr_, however, in the experiments we found the current to be highly variable in this window because of variations in access resistance among cells. We chose to focus on the 860-865 ms window, because it had the second largest I_Kr_ current isolation and was a few milliseconds after a voltage step, generating more consistent results. Figure 5C shows the change in current during the window from 860 ms to 865 ms for all cells in this study, separated by drug.

**Figure 5:**
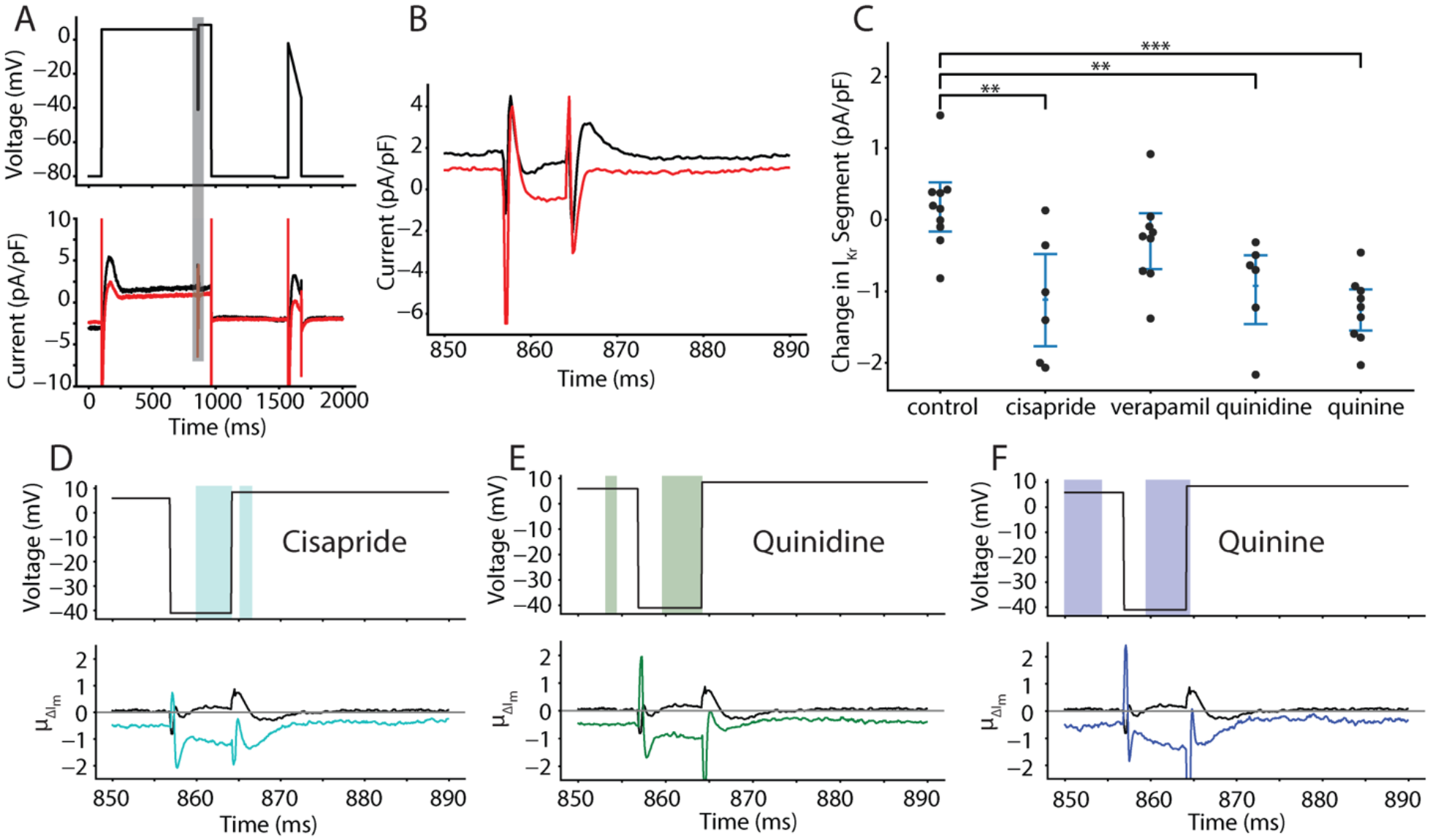
Optimized VC protocol correctly identifies I_Kr_ as a target of cisapride, quinidine, and quinine. **A**, Representative cell shows the effect of quinine on the current response during the segment of the VC protocol designed to isolated I_Kr_. The black trace is pre-drug and the red trace is post-drug. **B**, the current trace during the segment meant to isolate I_Kr_ (shaded grey in panel A). **C**, Cells treated with cisapride (p=.0032), quinidine (p=.0041), and quinine (p=.00002) show a decrease in total current during the VC segment designed to isolate I_Kr_. At the concentrations used in this experiment, cisapride, quinidine, and quinine should block 95%, 89%, and 72% of I_Kr_, respectively. **C**, **D**, and **E**, Functional t-tests show a significant difference in the average change in current during the I_Kr_-isolating segment when comparing cells treated with DMSO to cells treated with cisapride (**C**), quinine (**D**), and quinidine (**E**). Verapamil was excluded because there was no significant difference during this segment of the protocol.

Cells treated with cisapride, quinidine, and quinine all showed significant reductions during this segment compared to control cells. Cells treated with verapamil, a weaker I_Kr_ blocker at the concentration used here (Table I), did not show a significant difference in current during this segment. These data suggest that the VC protocol can detect strong I_Kr_ block during the model designed I_Kr_ segment but fails to detect light block of I_Kr_ current.

We followed this approach to identify the other major channels that were separately blocked by the protocol. The boxes marked with (*) in Table I indicate currents that were identified as being blocked based on significant changes in current during the corresponding model-identified segment. Importantly, all drugs that blocked more than 30% of a current were correctly identified, and there were no currents that were incorrectly identified as being blocked. This demonstrates that the VC protocol can be used to identify strong blocks of ionic currents with high sensitivity.

In addition to focusing on the model-identified segments of the protocol, we also performed a functional t-test where we calculated a p-value for the difference in current responses between DMSO- and drug-treated cells at every timepoint during the VC protocol. Figure 5C through 5E shows the segment of the VC protocol where I_Kr_ should be isolated. The colored window on the top of each panel shows the timepoints where there is a significant difference (p<.05) between control and drug treatment with cisapride (Figure 5D), quinidine (Figure 5E), and quinine (Figure 5F). The functional t-test identifies a significant difference in the currents between 860ms and 865ms for all three of these I_Kr_ blockers. There was no significant difference between DMSO and verapamil during this period.

The significance windows identified with a functional t-test were plotted over the entire VC protocol and suggest an agreement with the ionic currents that are blocked by each drug (Appendix – Figures 10-13). The verapamil-treated cells have a few brief windows of significance and most overlap with segments that the Kernik-Clancy model predicts will have I_CaL_ present. The cisapride- and quinidine-treated cells have longer and more significant windows. Most of the cisapride windows align with segments that the Kernik-Clancy model predicts will have I_Kr_. Most of the quinidine windows align with segments the Kernik-Clancy model predicts will consist of I_Kr_, I_to_, I_CaL_, or I_Ks_.

Taken with dynamic clamp AP data, these results indicate that the optimized VC protocol can be used to identify underlying currents responsible for changes in AP morphology. These ion channel targets can be further studied in expression cells to acquire dose-response data for specific drugs.

### The optimized VC protocol identifies a previously unreported quinine block of **I_f_**

Interestingly, our quinine-treated iPSC-CMs revealed a previously unreported block of I_f_. The protocol identified a significant reduction (p=.0097) in current during the portion of the protocol designed to isolate I_f_. The significance windows that fall after 4000 ms, which includes the portion of the protocol designed to isolated I_f_, align closely with Kernik-Clancy-predicted I_f_ (Figure 6A). Between 4000 ms and 6000 ms, when the membrane potential is hyperpolarized, there is loss of inward current after quinine treatment. During the second, long holding step between 4500 ms and 5500 ms, the difference between the pre-drug and post-drug traces increases and persists until the voltage is stepped to +50 mV. For the first ∼100 ms after this step, the traces flip, with the post-treatment trace showing a reduction in outward current before settling into a similar total current of ∼4.5 pA/pF at 6150 ms. These dynamics align closely with I_f_, as it is a positive current at positive holding potentials and will inactivate with a time constant around 40 ms when stepped to +50 mV. Figure 6C shows the significance windows (top), average change in current from pre-to post-drug application (middle), and Kernik-Clancy simulated I_f_ current (bottom). This figure suggests that there is a significant change in current throughout the period when the Kernik-Clancy I_f_ is present. These windows of significance largely agree with the model at both negative holding potentials, when I_f_ current should be negative, and positive holding potentials when the current should be positive. When we consider only the segment of the protocol where I_f_ current isolation is maximized, we find a significant increase in the current when compared to control cells (Figure 6D).

**Figure 6:**
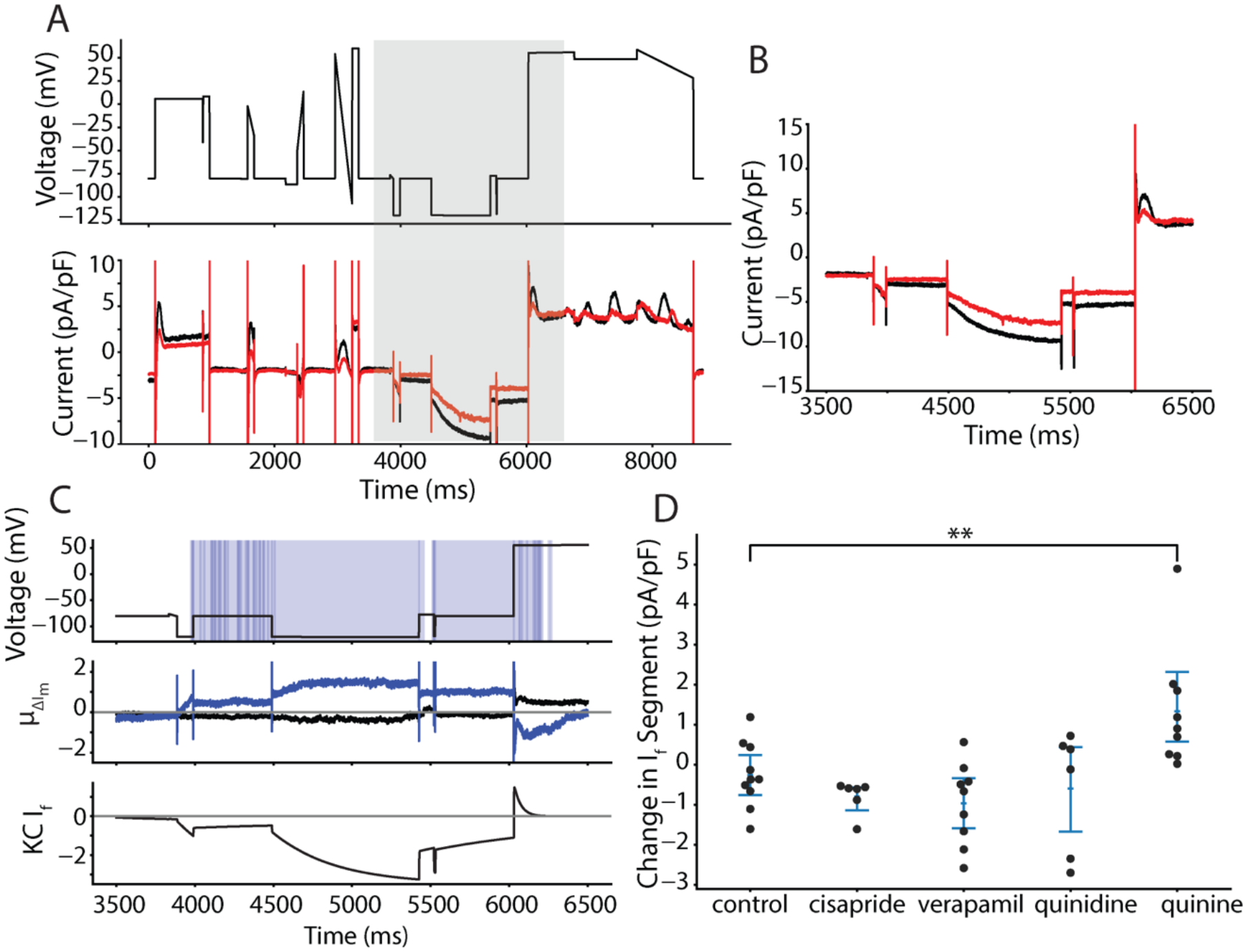
VC protocol identifies funny current as a novel target of quinine. **A**, Representative cell treated with quinine shows a reduction in inward current when the cell is clamped to a hyperpolarized potential of −120mV, before 6000ms. After 6000ms, outward current is reduced as I_f_ reverses in direction before inactivating. **B**, the current trace during the segment meant to isolate I_f_ (shaded grey in panel A). **C,** Functional t-test shows a significant difference between quinine-treated cells and control cells throughout the period of the protocol when funny current would be active (**C, top**). The average difference between quinine-treated and control cells is between 0.8 and 1.2 pA/pF throughout this period (**C, middle**). The Kernik-Clancy iPSC-CM model funny current becomes active during the period when quinine-treated cells show a significant change in current and turns off just before the p-value increases above .05 (**C, bottom**). **D**, Cells treated with quinine (p=.0097) show a decrease in total current during the VC segment designed to isolate I_f_.

To verify the finding that quinine blocks I_f_, we used a HEK-293 cell line stably expressing HCN1, the pore domain that conducts I_f_. The HCN1 isoform was chosen because it was recently shown to be present at high densities in iPSC-CMs (Giannetti et al., 2021). Figure 7A, B show that these HCN1 cells behave consistently with typical current-voltage I_f_ behavior. Dose-response data was acquired at seven concentrations of quinine (Figure 7C, D). The best fit Hill equation curve has an IC50 of 34.2 µM and Hill coefficient of 0.72. At 12 µM, which was the concentration used in the iPSC-CM study, the estimated block is 32.0%. Overall, this dose-response data confirms the findings from our iPSC-CM study, that quinine blocks I_f_ at 3x the EFPC.

**Figure 7:**
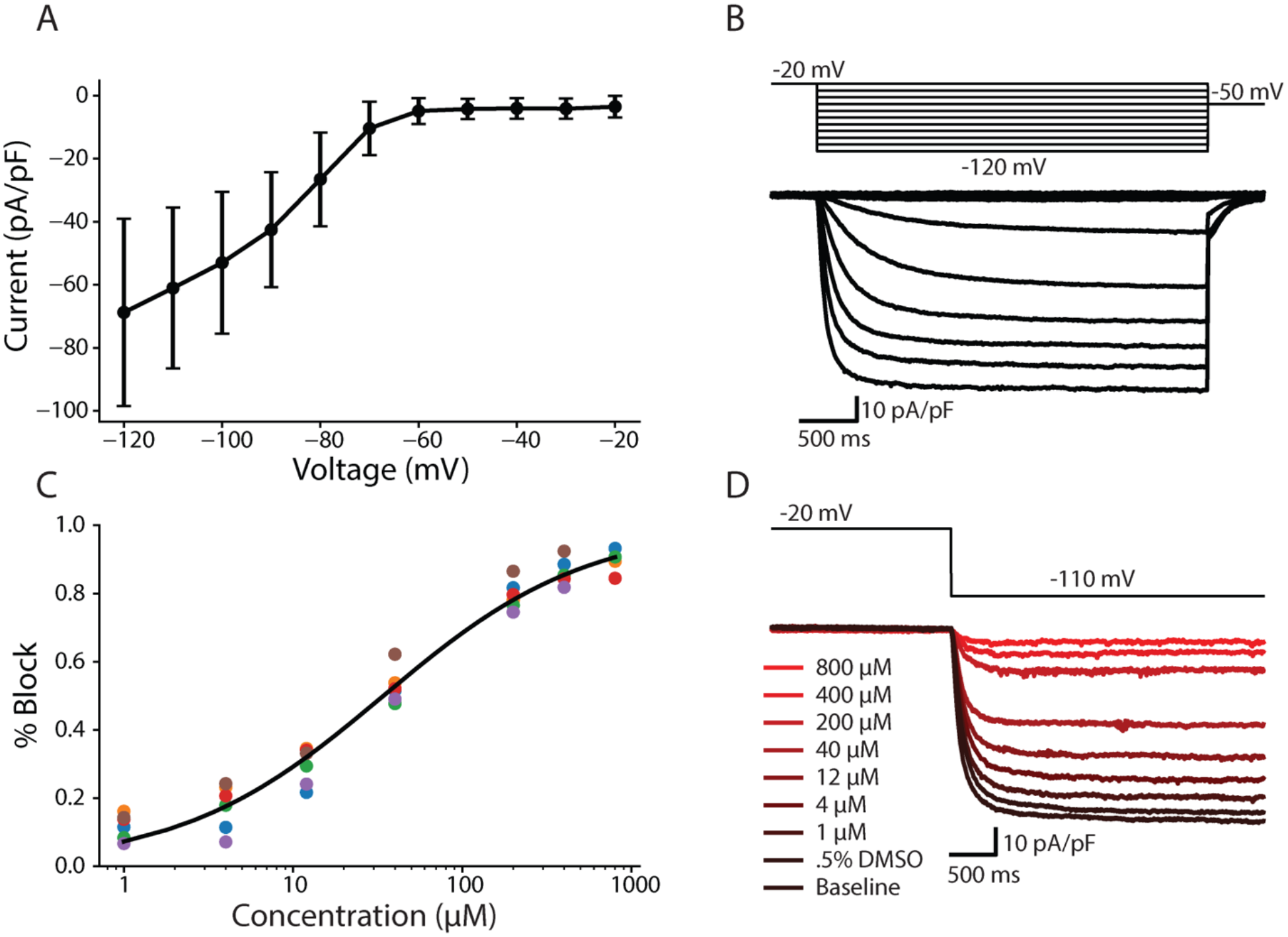
Dose-response curve shows quinine block of funny current in HEK-293 cells stably expressing HCN1. **A**, IV curve with averages and errors calculated from six HEK-HCN1 cells. **B,** Representative HEK-HCN1 current-voltage traces. **C,** Dose-response curve fit to pharmacology data from six HEK-HCN1 cells. **D,** Traces generated from a representative cell by clamping at −20mV for 3000ms, and then stepping to −110mV for 3500ms at all tested concentrations.

## DISCUSSION

In this study, we demonstrated the potential of a novel drug screening pipeline for predictive safety pharmacology. This approach simultaneously identifies surrogate markers of drug proarrhythmia in iPSC-CMs, and information about the drug’s underlying cardiotoxicity mechanism.

To overcome the difficulty of identifying cardiotoxicity markers and mechanism from individual iPSC-CMs, we developed an optimization algorithm that designed a VC protocol for the purpose of identifying multi-channel drug block. We then acquired optimized VC data, along with dynamic clamp AP data, from iPSC-CMs before and after drug treatment to identify markers of cardiotoxicity (e.g. AP prolongation) and mechanism (e.g. drug targets). This approach is the first of its kind to produce such detailed information about a drug’s action from individual iPSC-CM experiments. Using this approach, we also identified a novel block of I_f_ by quinine in iPSC-CMs. A similar such block was demonstrated in iPSC-derived neurons (Zou et al., 2018), but has never been confirmed in an expression system. We followed up on these findings by conducting a drug-response experiment using a cell line stably expressing HCN1 channels, and confirmed that quinine blocks HCN1 at the concentration used in this study.

### Optimizing VC protocols to improve experiments

Since Hodgkin and Huxley’s seminal work modeling the giant squid axon (Hodgkin and Huxley, 1952), there have been numerous efforts to fit electrophysiological models to less experimental data and to reproduce increasingly complex datasets. Traditional steady-state VC protocols take minutes to acquire and often result in models that poorly reproduce VC data from nonequilibrium states.

These shortcomings have been convincingly demonstrated with nonequilibrium response spectroscopy (Millonas and Hanck, 1998), which involves rapidly fluctuating voltage steps to quickly probe various states of an ion channel. These rapidly fluctuating, non-equilibrium protocols have been used to select the best Markov state model (Kargol, 2013; Kargol et al., 2004). Recently, a rapidly fluctuating 8-second VC protocol was developed and used to fit a hERG ion channel model (Beattie et al., 2018). This model was shown to outperform traditional steady-state-based models at reproducing validation data from over 5 minutes of recordings. These condensed VC protocols have made it possible to quickly acquire data under various external conditions and compare the results across multiple cells.

It has also been desirable to acquire rich VC data for fitting multi-channel models, such as neurons and cardiomyocytes. One such approach focused on quickly sampling the entire dynamic range of a neuronal cell, and using this data to fit model conductance values (Hobbs and Hooper, 2008). More recently, our lab developed a VC protocol that was manually designed to specifically isolate individual currents from a guinea pig cardiomyocyte to improve estimation of channel conductances (Groenendaal et al., 2015).

For both single- and multi-channel applications, most protocols have been designed with the ultimate goal of improving model fits. The current study is a departure from this approach in two ways: 1) the model used in this study was included as part of an optimization algorithm that designed the protocol, and 2) the resulting protocol was designed for the specific use of identifying drug targets, not for improving fits. In other words, the optimized protocol has a useful application outside of just improving cell model fits and predictions.

The success of this protocol in identifying ion channel targets is dependent on the quality of the underlying cell model (Kernik-Clancy) and inclusion of experimental artifact equations. Over the last couple decades, there has been an explosion in the number of human and animal cardiomyocyte models. With improvements in iPSC-CM maturity, health, and data quality there has been an increased interest in developing these models based on iPSC-CM data. These iPSC-CM models have been used to predict drug cardiotoxicity (Gong and Sobie, 2018; Jæger et al., 2021a; Tveito et al., 2018) and screen channel mutations (Kernik et al., 2020). Here, we used the model to design an optimal experiment rather than simulate experimental or clinical conditions. The success of the protocol in identifying drug cardiotoxicity targets may serve as a validation of the Kernik-Clancy and Paci iPSC-CM models. As mentioned above, we know that these *in silico* models can generate interesting hypotheses, simulate *in vitro* conditions, and guide therapy. However, to the best of our knowledge, this is the first work that shows how *in silico* models can be used to improve the design, and therefore impact, of cardiotoxicity drug studies.

### Genetic algorithm generates VC protocols that take advantage of the unique gating kinetics for each channel

The GA was designed to isolate individual currents, which resulted in protocols (Figure 2A) that take advantage of the unique gating kinetics for each ion channel. For example, the I_Kr_ protocol (Appendix – Figure 5) takes advantage of the fast-inactivation, slow-activation gating that is characteristic of this channel. The initial step to just above 0 mV quickly inactivates the channel. Over the course of a few hundred milliseconds, the activation gate opens. Little current flows through the channel at this voltage because the inactivation gate is almost entirely closed. The step to −40 mV opens the inactivation gate, allowing current to flow through the channel before the slow activation gate closes. The step back above 0 mV increases the driving force, which provides a brief window (about 5 ms) where the activation gate is open, the inactivation gate is open, and there is a large driving force pushing potassium into the cell.

The other protocols also identified dynamic ranges that highlight each channel’s unique kinetics. The I_Ks_ (Appendix – Figure 7) and I_f_ (Appendix – Figure 8) protocols settle into positive (>50 mV) and negative (<-110 mV) extremes where most other channels are closed, but these channels are open and remain open. The I_K1_ protocol (Appendix – Figure 6) isolates its current at the same potential as I_f_, but is maximized before I_f_ has a chance to open. The I_Na_ protocol (Appendix – Figure 2) steps to a hyperpolarized potential (−87 mV) to open the activation gate and then jumps to a ramp that is depolarized enough to activate the channel (−50 mV), while minimizing the activation of other ion channels (e.g. I_CaL_). The I_CaL_ and I_to_ channels have fast activation and slow inactivation kinetics – these protocols take advantage of this by stepping to potentials that will open their channels but minimize the contribution from the other.

### Drug cardiotoxicity screening

The current, overly sensitive cardiotoxicity screening guidelines points to the need to develop new methods that improves specificity and provides insight into the mechanism of drug block. In recent years, high throughput iPSC-CM screening approaches that rely on surrogate markers of cardiotoxicity risk have improved the ability to evaluate drugs at scale and with improved accuracy compared to traditional methods (Bedut et al., 2016; Lu et al., 2019; Pioner et al., 2019). Expression system cell lines and molecular dynamic simulations have supplemented these findings by providing detailed mechanisms of drug action at the single-channel level (Demarco et al., 2020; Yang et al., 2020). However, there have been few methods that provide both measures of cardiotoxicity and mechanism from the same iPSC-CM cells.

One recent approach to address this need was to fit an iPSC-CM model to fluorescent voltage and calcium AP data acquired before and after drug application (Jæger et al., 2021a, 2021b; Tveito et al., 2018). With this method I_Kr_, I_CaL_, and I_Na_ percent block was determined. Because this approach only considers spontaneous AP and conduction velocity data, successful estimates of current block is limited to the currents (e.g. I_Kr_, I_CaL_, and I_Na_) that are sensitive to changes in these data.

In the current study, we build upon this work by developing a pipeline that provides both surrogate measures of drug cardiotoxicity and identification of drug block for seven cardiac ion channels. With the HEK-293 study, we showed the potential of using expression cell lines to confirm findings in iPSC-CMs and determine potency at multiple concentrations.

### Limitations and Future Directions

This study has some limitations. First, before this approach can be used at scale, it must first be validated with a high-throughput automated patch-clamp system. With this type of system, data acquisition could be 10-100x faster and operators would not need the specialized patch-clamp skill. Because the automated patch-clamp system applies the same protocol identically every time, these studies will likely have more consistent experimental artifact parameters, such as leak and access resistance (Goversen et al., 2018a), and increased power for statistical analyses. Also, the microfluidic administration of drugs in these systems allows for quick wash-on steps, making it possible to acquire data at more concentrations. This would provide dose response data in each cell, which could be used to identify the relative size of the currents present.

Second, in addition to improvements made possible by an automated system, some adjustments to the VC protocol and data analysis could improve the mechanistic insights made in future studies. Additional channels besides the seven considered in this study could be included. This is easy to address, by simply adding currents to the VC protocol optimization algorithm. One challenge that would require additional adjustments is to design an algorithm that can tease apart drug effects on ion channel kinetics. An optimization algorithm to address ion channel kinetics would need a target objective that is, likely, very different from the one used in this study.

Third, when treated with a high dose of cisapride, the iPSC-CMs in this study did not show significant APD_90_ prolongation, despite its strong block of I_Kr_. This may be due to the cells having a relatively low density of I_Kr_ compared to adult cardiomyocytes. Quinidine and quinine, also strong I_Kr_ blockers, did cause significant prolongation. This may have been caused by their small block of I_Ks_, which would further deplete the repolarization reserve of these cells. Further tests with cisapride would be needed to determine whether this was an issue of statistical power.

Fourth, the iPSC-CMs often produced an oscillatory current trace when held at large positive voltages (e.g. from 6500 to 9000 ms in the VC protocol). This is likely caused by calcium overload that can occur at high potentials. This could decrease the sensitivity of the protocol to determine strong I_Ks_-blockers. In the future, we would like to test the ability of the protocol to strong blocks of I_Ks_.

### Conclusion

In this study, we outline a new pipeline for determining drug cardiotoxicity and underlying mechanism by applying a novel VC protocol and I_K1_ dynamic clamp to iPSC-CMs. By analyzing changes in AP and VC data acquired after drug application, we were able to identify cardiotoxicity markers and currents that were strongly blocked by the drug. We also identified a novel block of I_f_ by quinine, which was confirmed using an expression system cell line. In the future, the scalability of this method can be improved with an automated patch clamp system and the detail of the mechanistic insights can be increased by applying this protocol to iPSC-CMs at multiple drug concentrations. We think that this cardiotoxicity pipeline could have far-reaching effects on how drugs are screened and could ultimately increase the number of safe and effective drugs available to patients.

## MATERIALS AND METHODS

### iPSC-CM and artifact model

The baseline Kernik-Clancy iPSC-CM model was used in this study (Kernik et al., 2019). Prior to the VC protocol optimization, the model was run to steady state, and then simulated under voltage clamp at −80 mV for 20 s. We included the simplified experimental artifact equations from Lei et al. in our model simulations (Lei et al., 2020). In this recent study, these artifact equations were incorporated into an ion channel model and shown to produce better fits to experimental data, and with fewer parameters. The effects of these patch clamp artifacts are particularly pronounced within the first few milliseconds after a voltage step (Appendix – Figure 1). This was an important factor to consider in our optimization because we anticipated the optimal protocols may isolate currents during these windows. This is based on the observation that many existing VC protocols designed to maximize current through a single ion channel (e.g hERG or Na_V_1.5) often do so within the first few milliseconds after a voltage step, where artifacts most obscure current readings.

The following equations summarize the iPSC-CM model and artifact equations used in our simulations:

The I_ion_ equation is the sum of all ionic currents from the Kernik-Clancy iPSC-CM model. The leak current equation quantifies the amount of contamination caused by an imperfect seal between the pipette tip and cell membrane. The observed current is a sum of the leak and ionic currents. The series resistance compensation equation affects the speed with which the cell membrane voltage will reach the command voltage after a step. The membrane voltage differential includes a term for the voltage offset. This term is a catchall offset from the amplifier, electrode, and liquid junction potential. Because we zero the amplifier offset before patching each cell, we assume that this term is equal to our experimental liquid-junction potential (−2.8 mV) in our simulations. The Kernik-Clancy model with experimental artifact was implemented using custom Python code. Below is a list of parameter definitions, along with the values used in our model simulations.

### Voltage clamp protocol optimization

An optimized VC protocol was designed for each of seven currents (I_Kr_, I_CaL_, I_Na_, I_to_, I_K1_, I_f_, and I_Ks_). To optimize the protocol, we used custom Python code that implemented a genetic algorithm (GA) with the DEAP Python package (Fortin et al., 2012). The GA had 200 individual protocols per generation and 50 generations. Below, we discuss how we implemented the initialization, evaluation (i.e. calculated cost), selection, mating, and mutation functions of the GA.

#### Initialization

Each individual in the GA was initialized with a random set of four voltage segments. Each segment could be either a step or ramp between 5 and 1000 ms long and with voltages between −120 and 60 mV. If the segment was a step, the duration was randomly selected between 5 and 1000 ms and the voltage was randomly selected between −120 and 60 mV. If the segment was a ramp, the duration was randomly selected between 5 and 1000 ms, and the start and end voltages were selected between −120 and 60 mV.

#### Evaluation

To evaluate the fitness of a VC protocol, we clamped the Kernik-Clancy model, and calculated the percent contribution (C(t)) of the target current at every timepoint during the protocol.

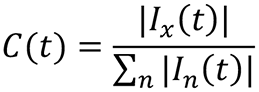

In this equation, I_x_ is the target current and I_n_ is any current that contributes to the observed current (I_out_). The denominator is the sum of the absolute values for all these currents, including ionic currents and the artifact leak current. The best possible contribution score is one and represents when the target current contributes all the observed current. The worst contribution score is zero and represents when the target current is turned off. We calculated the average contribution, C(t), over a 10 ms window at each timepoint and used the highest contribution window value as the fitness score for the protocols.

#### Selection

To select the protocols that continue to the next generation, we used k=2 tournament selection with replacement. This means that the GA ran 200 random head-to-head matchups, and the protocol with the higher fitness score in each matchup moved into the mating pool. Because we randomly selected protocols from the same pool for every matchup, it was possible for some individuals to be in multiple matchups, while others were not in any matchups. After all the matchups, there were 200 individuals in the mating pool. It was common for this new pool to include multiple copies of high-fitness protocols.

#### Mating

Each individual in the mating pool was paired with one other individual and uniform crossover was applied to create two child individuals. For each pair, there was a 90% chance that the individuals would mate. If the individuals did not mate, the protocols would remain the same and their features would be cloned into two new individuals and placed in the mutation pool. If individuals did mate, there was a 20% chance of swapping each segment in their protocol. The offspring protocols that resulted from the swapping of segments between the two individuals were moved into the mutation pool.

#### Mutation

For each protocol in the mutation pool, there was a 90% chance of being selected for mutation. For each segment in a selected protocol, there was a 10% chance of mutation. To mutate the duration of a segment, a random number was selected from a normal distribution centered around zero, with a standard deviation of 47.8 ms, and added to the existing duration. To mutate the voltage of a segment, a random number was selected from a normal distribution centered around zero, with a standard deviation of 6 mV, and added to the existing voltage. If the resulting duration or voltage was outside the bounds (e.g. 5 to 1000 ms or −120 to 60 mV), a new mutation value would be selected until the duration or voltage were valid. Once all protocols were considered for mutation, the population was moved to the next generation where selection begins again.

Evaluation, selection, mating, and mutation were repeated for 50 generations.

#### Combining protocols

We took the protocol with the highest current contribution for each of the seven ionic currents (Supp Figs II-VIII) and combined them into one large protocol. The durations of the seven protocols ranged from ∼1400 ms for I_to_ to ∼3800 ms for I_Ks_ (Figure 2A). Before combining the protocols, we systematically shortened them using a two-step process. First, we removed the portion of the protocol more than 50 ms after the maximum current contribution window. Then, we incrementally removed 10 ms segments from the beginning of the protocol, while ensuring the max current contribution did not decrease by more than 5%. The seven shortened protocols were then connected by 500 ms holding steps at −80 mV. We chose to hold at −80 mV for 500 ms, because all of the protocols were optimized after a −80 mV holding step, and because this is long enough for the kinetics of most ion channels to reach steady state.

To validate that the VC protocol isolates current during the same time windows in a cell with different conductance and kinetic parameters, we applied the optimized VC protocol to the Paci iPSC-CM model (Paci et al., 2018) with experimental artifacts. The Paci model, which was originally designed based on different data from the Kernik-Clancy model, can be interpreted as a cell with different kinetic and conductance parameters. This was an important step because of the heterogeneity among iPSC-CMs.

We found the time windows that maximized the current isolation for each of the seven currents using the the Paci-artifact model and compared those time windows to the Kernik-Clancy-artifact results (Appendix – Figure 9). Five (I_K1_, I_to_, I_Kr_, I_Ks_, and I_Na_) of the seven currents in the Paci model were isolated within 10 ms of the Kernik-Clancy model. The maximum I_CaL_ isolation in the Paci model occurs far from where the current is maximized in the Kernik-Clancy model. However, the Paci model had a current isolation within 5% of its maximum during the Kernik-Clancy window. The timepoint for I_f_ also differed between the two models. However, these timepoints are near one another and have similar voltage dynamics. These data provide a signal that the Kernik-Clancy-designed protocol is generalizable enough to isolate currents in iPSC-CMs with different kinetics and conductances.

All code for this pipeline has been made available on the Christini Lab GitHub: https://github.com/Christini-Lab/vc-optimization-cardiotoxicity.

### Human iPSC-CM experiments

#### iPSC-CM cell culture

Frozen stocks of Human induced pluripotent stem cell-derived cardiomyocytes (hiPSC-CMs) from a healthy individual (*SCVI-480CM*) were obtained from Joseph C. Wu, MD, PhD at the Stanford Cardiovascular Institute Biobank. All iPSC-CM lines obtained from the individual were approved by Stanford University Human Subjects Research Institutional Review Board and differentiated to cardiomyocytes as described previously (Burridge et al., 2014; Churko et al., 2013).

Each vial of iPSC-CMs was cultured as a monolayer in one well of a 6-well plate precoated with 1% Matrigel and supplemented with RPMI media *(Fisher/Corning 10-040-CM*) containing 5% FBS (*Gibco 16000069*) and 2% B27 (*Gibco A1895601*). Cells were placed in an incubator at 37°C, 5% CO_2_, and 85% humidity for 48 hours. When replating, cells were lifted with 1 mL Accutase (*Corning A6964*), and the enzymatic reaction was blocked with DMEM/F12 (*Gibco 10565-042*) plus 5% FBS (Burridge et al., 2014). Cells were diluted to 100,000 cells/mL and re-distributed to 124 8 mm sterile coverslips precoated with 1% Matrigel. RPMI media was replaced every other day. Cells were patched from days 5 to 15 after thaw.

#### Electrophysiological Setup

Borosilicate glass pipettes were pulled to a resistance of 2-4 MΩ using a flaming/brown micropipette puller (Model P-1000; Sutter Instrument, Novato, CA). The pipettes were filled with intracellular solution containing 10 mM NaCl, 130 mM KCl, 1 mM MgCl_2_, 10 mM CaCl_2_, 5.5 mM dextrose, 10 mM HEPES. Amphotericin B was used to perform perforated patch. The pipette tip was first dipped into intracellular solution with no amphotericin B for 2-5 s. The pipette was backfilled with the intracellular solution containing 0.44 mM amphotericin B. The coverslips containing iPSC-CMs were placed in the bath and constantly perfused with an extracellular solution at 35-37°C containing 137 mM NaCl, 5.4 mM KCl, 1 mM MgSO_4_, 2 mM CaCl_2_, 10 mM dextrose, 10 mM HEPES.

Patch-clamp measurements were made at 10kHz by a patch-clamp amplifier (Model 2400; A-M Systems, Sequim, WA) controlled by the Real Time eXperiment Interface (RTXI; http://rtxi.org) to send commands to the amplifier via the data acquisition card (PCI-6025E; National Instruments, Austin, TX). After immersing the pipette into the extracellular solution, voltage was set to zero, and voltage offset in our recordings was assumed to be equal to the liquid junction potential of −2.8 mV. After contact with a cell was made and a seal of greater than 300 MΩ was established, we waited for the access resistance to decrease below 40 MΩ before starting experiments. The series resistance was 9-40 MΩ for all experiments, and series resistance compensation was set to 70%. The 70% compensation was chosen because larger values caused oscillations during the recordings.

#### Experimental design and drugs

Spontaneous, I_K1_ dynamic clamp, and VC data was acquired before and after drug application. Once access was gained, spontaneous behavior was acquired for >10 s. Dynamic clamp I_K1_ data was acquired using a custom RTXI module that implemented the Ishihara et al. (2009) model. A recent *in* silico study showed that the Ishihara model has properties that are optimal for use in I_K1_ dynamic clamp studies of hiPSC-CMs (Fabbri et al., 2019). With this module, we incrementally increased the Ishihara I_K1_ conductance by 0.25x of its baseline conductance until spontaneous behavior stopped, and the cell reached a resting membrane potential below −65 mV. Resting at this hyperpolarized potential allows recovery of sodium channels, resulting in APs with faster upstrokes and larger amplitudes, better resembling adult ventricular APs. All but one cell settled into a resting membrane potential below −69 mV (Figure 3C). After dynamic clamp data was acquired, the amplifier was switched to voltage clamp mode, and compensation of capacitance and access resistance was done. The cell was then clamped with the optimized VC protocol.

Following VC acquisition, the perfusion system was switched to an external solution containing either 0.1% of DMSO, or one of the following drugs: cisapride monohydrate at 250 nM (USP – SKU: 1134120, Rockville, MD), verapamil hydrochloride at 150 nM (MP Biomedicals – SKU: 195545, Solon, OH), quinidine at 2.7 µM (Tocris – SKU: 4108/50, Bristol, UK), or quinine at 12 µM (Sigma-Aldrich – SKU: 22620, Saint Louis, MO). Drug solutions were prepared daily, by dissolving in DMSO before addition to external solution. The DMSO concentration was <0.1% for all drug solutions. Cells were exposed to the drug solution for >5 minutes, while square pulses were applied to observe changes in the current response. Once the cell had been exposed for >5 minutes, and changes in the current response had stabilized, spontaneous, I_K1_ dynamic clamp, and VC data was acquired by following the same steps above.

### Data Analysis and Statistics

All results are presented as mean ± standard error of the mean. Significant differences between the DMSO and each drug group were calculated using the SciPy unpaired t-test function in Python, with significance indicating p<0.05. The precise p-value for each statistical test is presented in its corresponding figure. Confidence intervals are set to 95% for each point plot. All statistical analyses were performed using the raw experimental data. For presentation in figures, data were smoothed with a 0.4 ms moving average.

A power analysis was not used to make a sample-size estimation because we saw significant differences between groups after experiments conducted on one freshly thawed batch of cells for each drug.

#### Analyzing AP features

The resting membrane potential (RMP), action potential amplitude (APA), action potential duration at 20% repolarization (APD_20_), and action potential duration at 90% repolarization (APD_90_) were calculated for all dynamically clamped I_K1_ studies using custom Python code. The RMP was determined by finding the minimum voltage during an AP. The APA was calculated as the difference between the RMP and the maximum voltage during an AP. The APD_20_ and APD_90_ were calculated as the duration between the maximum upstroke velocity timepoint and when the cell repolarized to 20% and 90% of its RMP.

#### Analyzing VC protocol data

##### Functional t-test

A functional t-test (Keser, 2014) was used to determine the time windows when current response changes to drug treatment differed from responses to DMSO treatment. We used the following steps to develop a null distribution, conduct a functional t-test, and determine windows of significant difference between DMSO control and drug groups:

1. For both the control and drug groups, we calculated the change in current at every timepoint from pre-to post-treatment.
2. We developed a null distribution by completing the following step 200 times: we combined and randomly shuffled individuals from the drug and control groups, and then redistributed them into two distinct groups. We calculated a T-statistic (T(t)) at every timepoint.
3. We found the t-value at the 95^th^ quantile of the null distribution and used it as the threshold for determining significant differences between our control and drug groups. In other words, the control and drug groups were labeled significantly different at a time when the T-value comparing these two groups was greater than the T-value at the 95^th^ percentile of the null distribution.

The windows plotted in Figures 5, 6, and Appendix – Figures 10-13 show where there is a significant difference between the drug and DMSO groups that lasts for more than 1 ms. The functional t-test calculations were completed with custom Python code using the SciPy unpaired t-test function.

### HEK-HCN1 culture

Human embryonic kidney cells 293 stably expressing human hyperpolarization-gated cyclic nucleotide-sensitive cation channel 1 (HEK-HCN1) were obtained from Charles River *(CT6114)*. Cells were cultured and maintained according to the online protocol by Charles River. One frozen vial of cells was thawed in prewarm DMEM/F12 media plus 10% FBS and 100 units/ml Penicillin/Streptomycin *(Life Technologies 15140)* and placed in 37°C, 5% CO_2_, and 85% humidity incubator overnight. The media was replaced with selection media containing 0.005 mg/mL Blasticidin *(InvivoGen ant-bl-5b)* and 0.1 mg/mL Zeocin *(InvivoGen ant-zn-5b),* and the cells were sub-cultured if they were at ∼ 75% confluency.

To induce expression of HCN1 channels, cells were cultured in DMEM/F12 media plus 1.5 µg/mL tetracycline *(Sigma-Aldrich T7660)* two days before the experiment. To prepare the cells for use in the Nanion Patchliner automated patch system, they were rinsed twice with 5 mL Hank’s Balanced Salt solution *(Life Technologies 14175)* and lifted with 2 mL Accutase. The enzymatic reaction was blocked using DMEM/F12 media and the cell solution was centrifuged for 2 min. Supernatant was discarded and the cell pellet was mixed with FBS-free DMEM/F12 media plus 15 mM HEPES (pH 7.3) and extracellular solution. The cell mix solution was placed in a 4°C fridge for 10 min before use.

### HEK-HCN1 Experiments

HEK-HCN1 current was recorded using an automated patch clamp system (Patchliner Quattro, Nanion Technologies GmbH), sampling at 25kHz. Whole cell patch clamp was performed at room temperature on HEK-HCN1 using standard medium resistance NPC-16 chips (1.8-3 MΩ) after getting GΩ seal. Series resistance was compensated at 80% for voltage-clamp recordings.

I_f_ Current-voltage (I-V) traces were recorded starting from a holding potential of −20mV. Cells were stepped to a potential between −20 mV and −120 mV for 3500 ms, decreasing by 10 mV steps. Then, cells were stepped to −50 mV for 500 ms to acquire the tail current. Maximum currents were calculated as the average current over the last 800 ms of the hyperpolarizing step. Maximum tail currents were calculated as the average between 8 and 16 ms after the step to −50 mV.

To measure the dose-response of HCN1 current to quinine treatment, peak currents were measured by stepping from a holding potential of −20 mV to −110 mV for 3500 ms. Maximum currents were calculated as the average current over the last 800 ms of the hyperpolarizing step. These traces were acquired three or four times at each dose. The last two traces at each dose, which were acquired >40 s after drug application, were used for analysis. Each data point plotted in Figure 7C is an average of the currents from these two traces.

In all experiment, cells were measured using extracellular solution with the following concentrations (in mM): 140 NaCl, 4 KCl, 2 CaCl2, 1 MgCl2, 5 D-Glucose monohydrate, 10 HEPES, pH adjusted to 7.4 with NaOH, and 298 mOsm. The intracellular solution had the following concentrations (in mM): 10 EGTA, 10 HEPES, 10 KCl, 10 NaCl, 110 KF, pH 7.2 with KOH, 280 mOsm.

### Code and data availability

All code has been made publicly available on GitHub at: https://github.com/Christini-Lab/vc-optimization-cardiotoxicity. Data that support the findings can be provided upon request.

## ACKNOWLEDGMENTS

We thank Dr. Drew Tilley for his experimental insights and expertise. We thank Dr. Sumanta Basu for consulting on the functional t-test and Dr. Henry Sutanto for feedback on the manuscript.

## FUNDING

Research reported in this publication was supported by the National Heart, Lung, And Blood Institute of the National Institutes of Health under Award Number F31HL154655 (to A.C.) and U01HL136297 (to D.J.C.). The content is solely the responsibility of the authors and does not necessarily represent the official views of the National Institutes of Health.

## APPENDIX

**Appendix – Figure 1.**
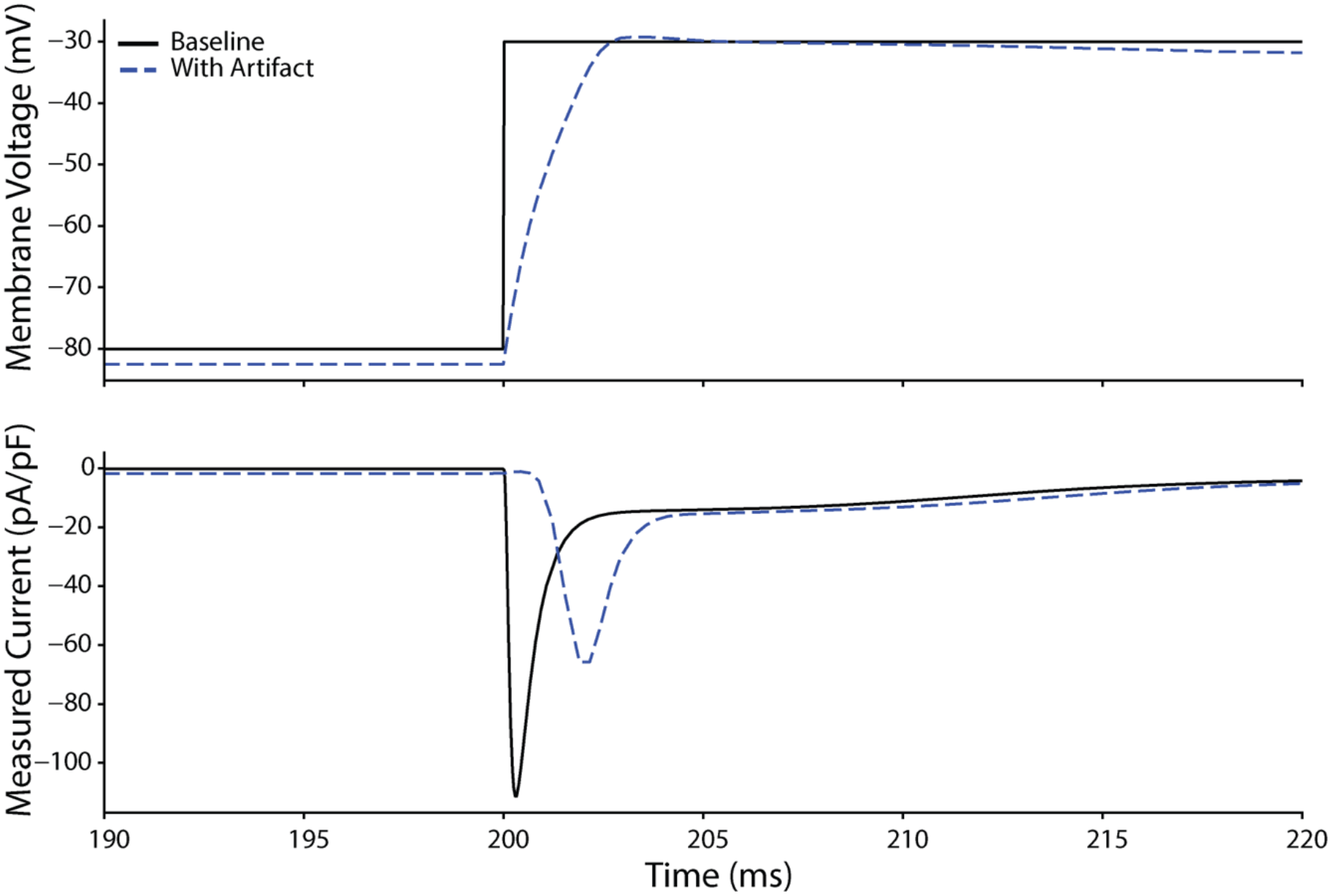
The effect of experimental artifact on voltage clamp data designed to activate sodium channels. The experimental artifact used in this simulation included a voltage offset of −2.8 mV, seal resistance of 1 GΩ, and access resistance of 20 MΩ. The top panel shows the voltage experienced by the cell (dashed blue) compared to the command voltage (black). The voltage offset shifts the membrane voltage negative by 2.8 mV, which has little effect on the current response. The relatively high access resistance is what causes the gradual slope upwards from the starting voltage of −80 mV to the ending voltage of −30 mV. This gradual slope in the membrane voltage leads to a delayed and reduced peak current (bottom) response.

**Appendix – Figure 2.**
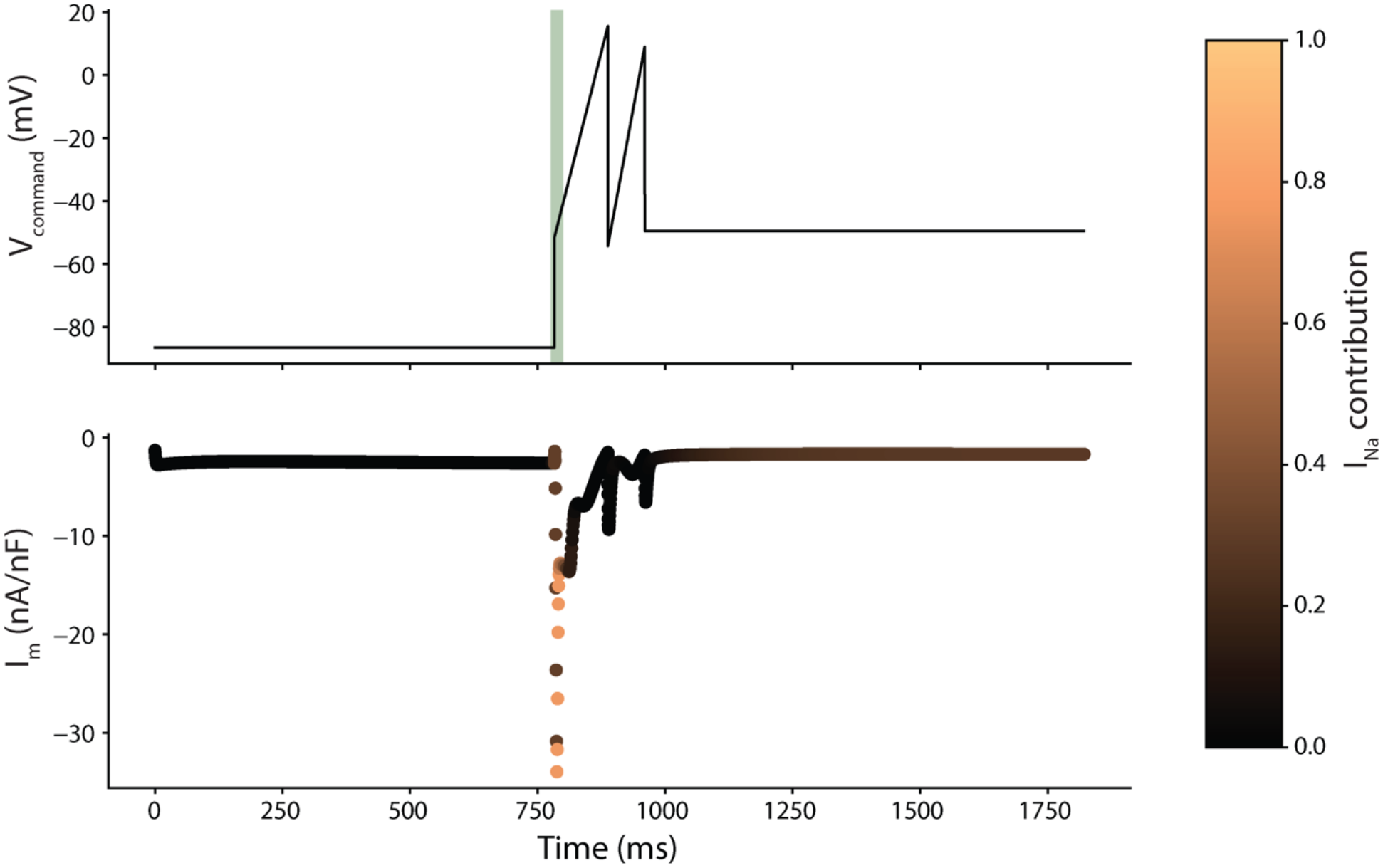
The optimized protocol for I_Na_ with Kernik-Clancy current response. The optimized protocol for I_Na_ is shown in the top panel. The bottom of panel shows the Kernik-Clancy response to the protocol. Each point plotted in the bottom panel is color-coded by the relative contribution of the specified current. The green line in the top of each panel shows where the isolated current is maximized.

**Appendix – Figure 3.**
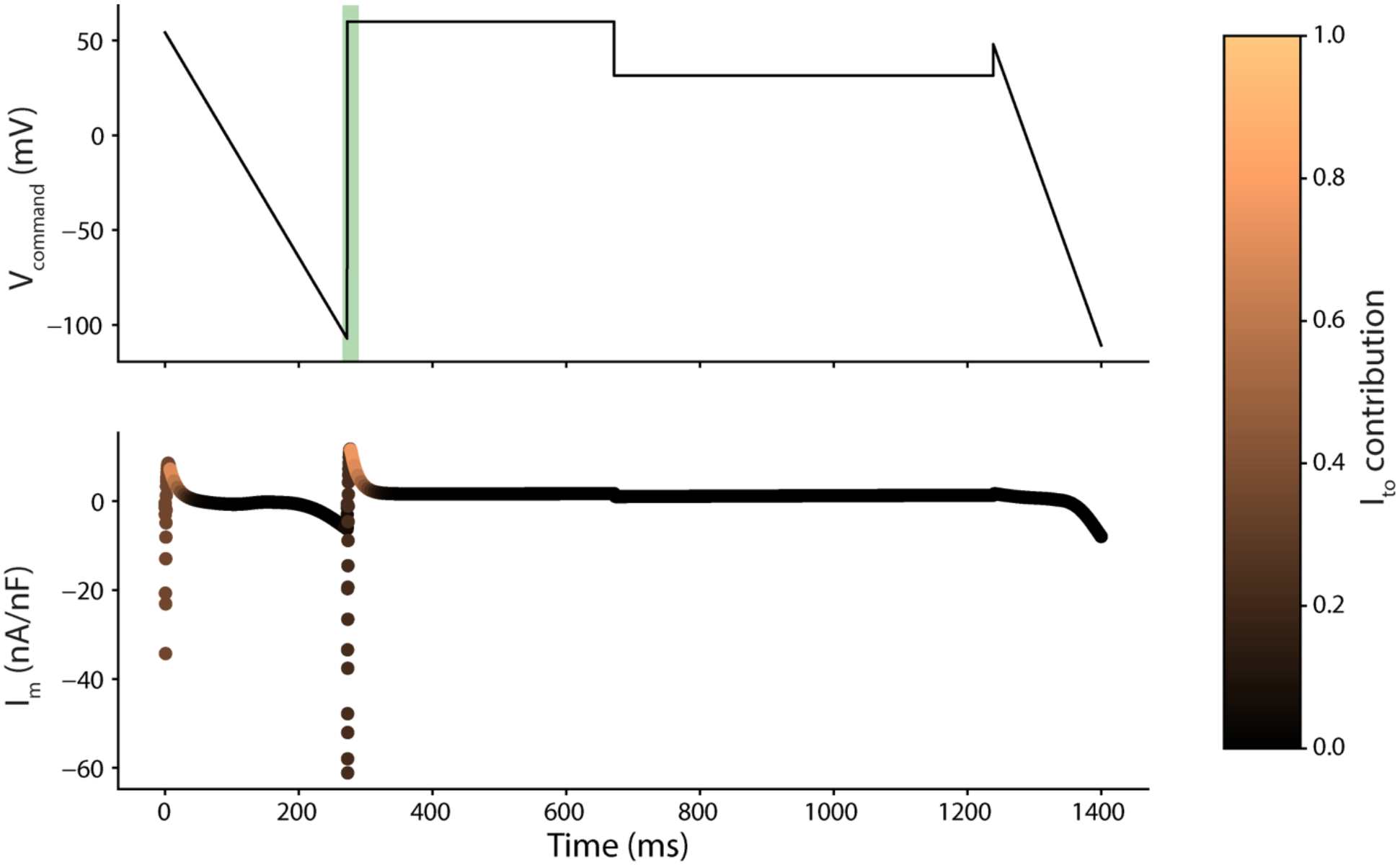
The optimized protocol for I_to_ with Kernik-Clancy current response.

**Appendix – Figure 4.**
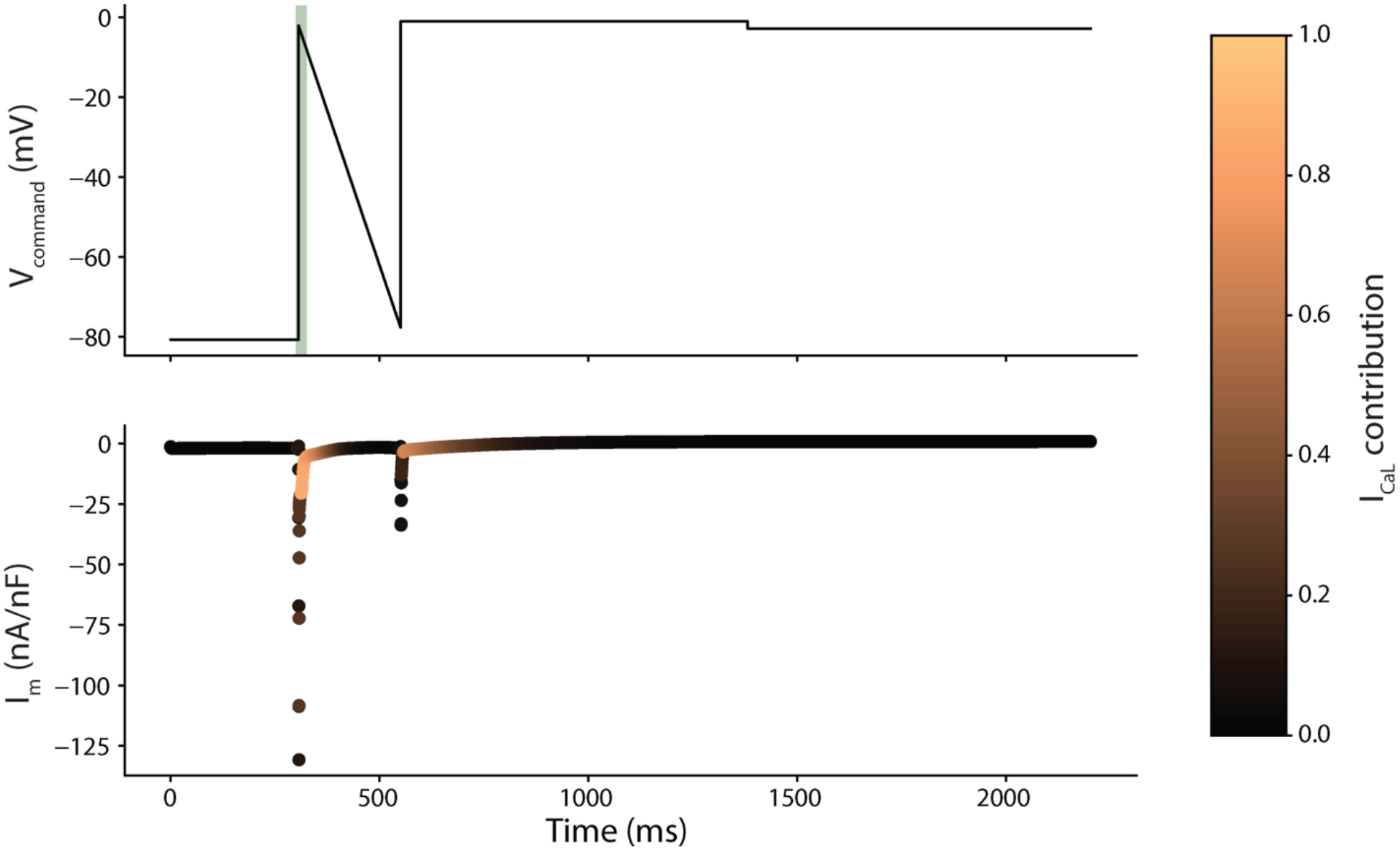
The optimized protocol for I_CaL_ with Kernik-Clancy current response.

**Appendix – Figure 5.**
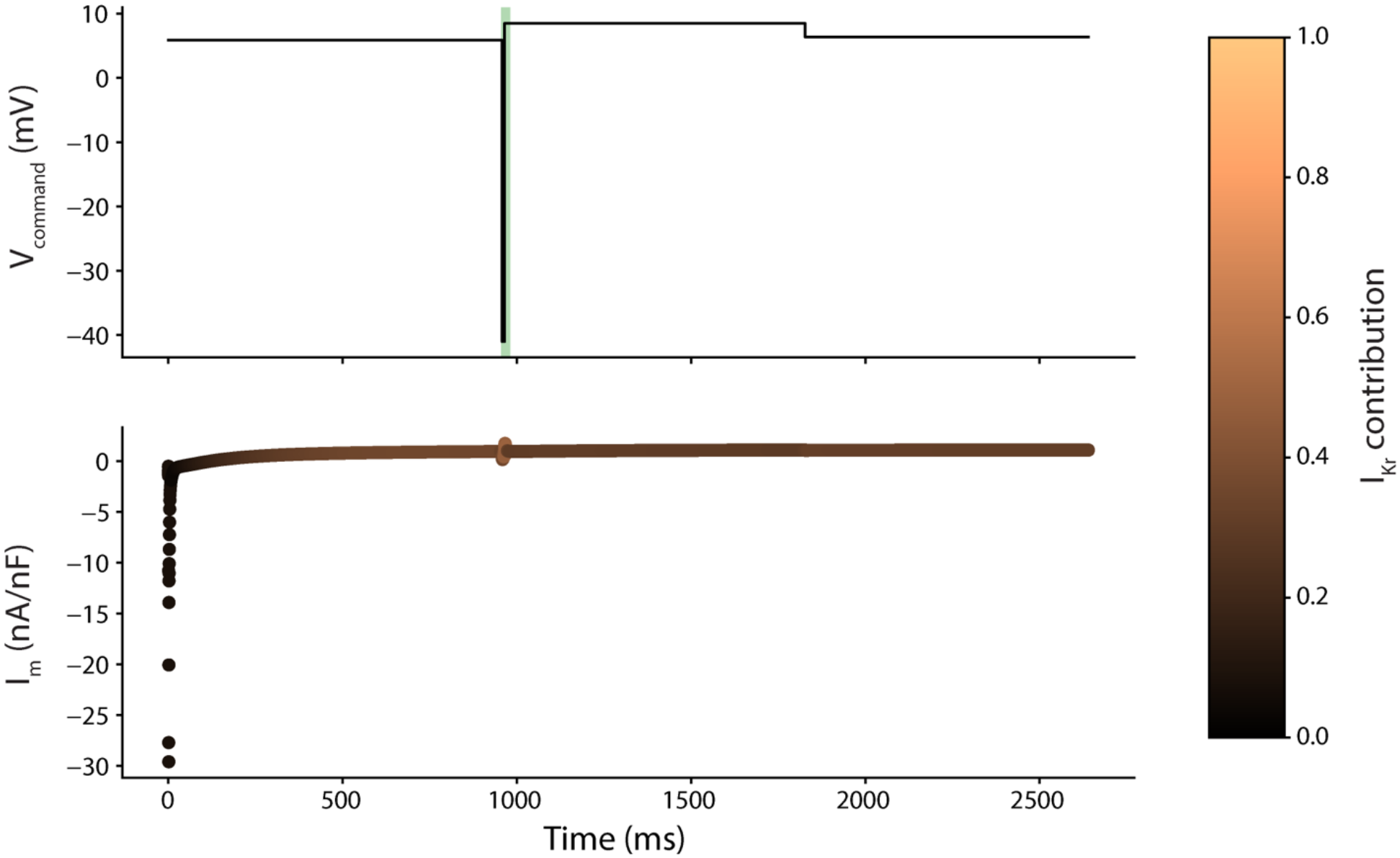
The optimized protocol for I_Kr_ with Kernik-Clancy current response.

**Appendix – Figure 6.**
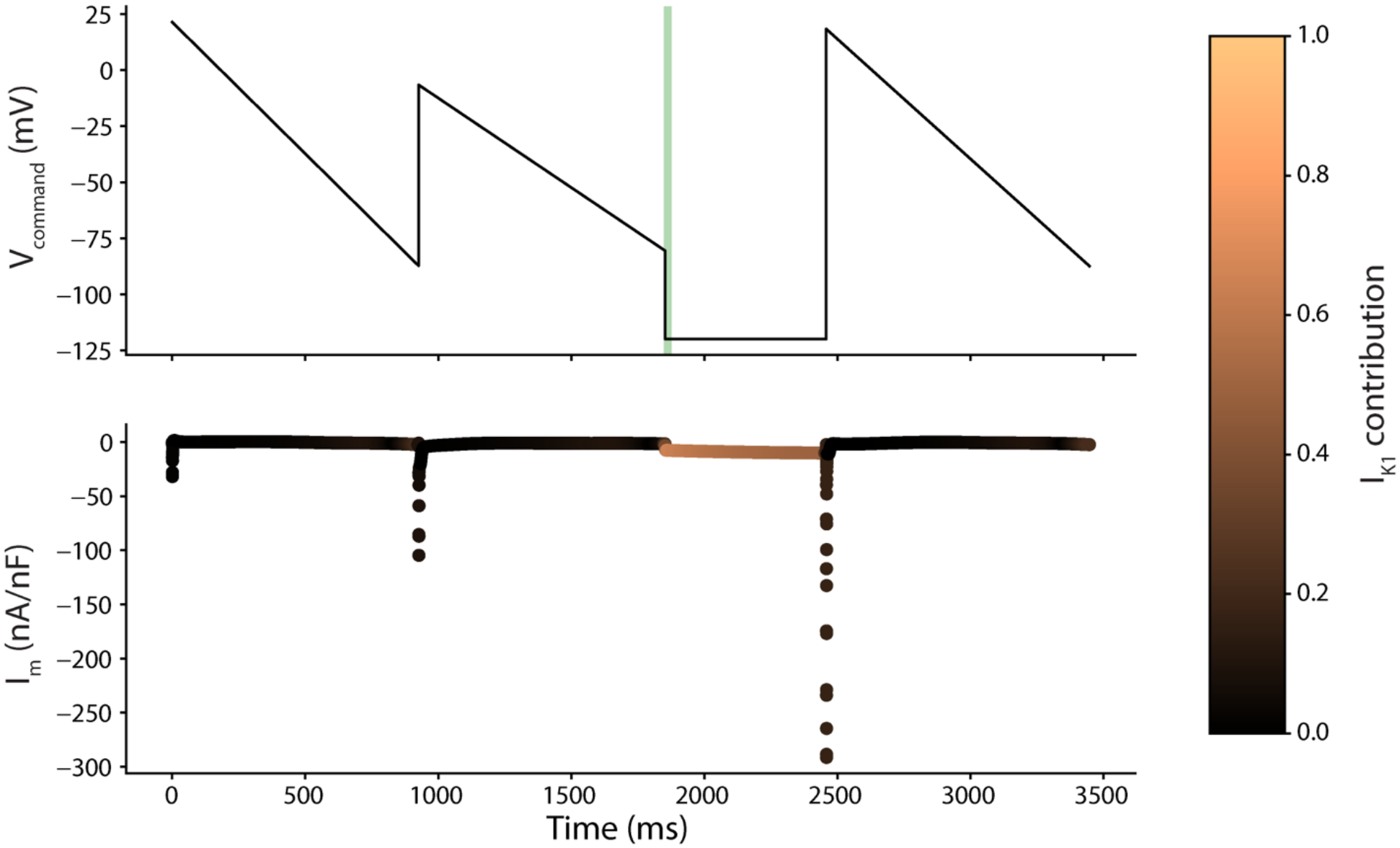
The optimized protocol for I_K1_ with Kernik-Clancy current response.

**Appendix – Figure 7.**
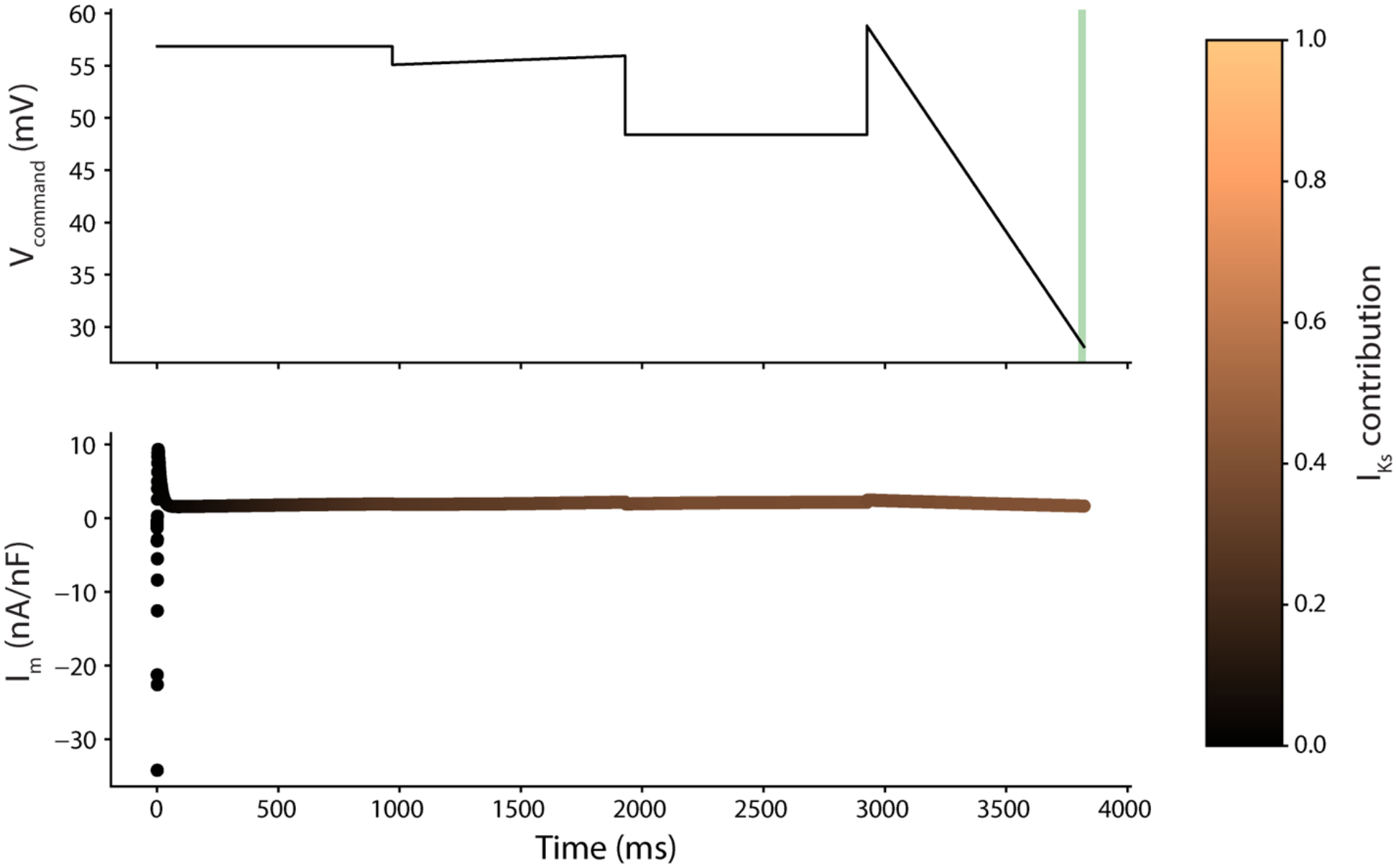
The optimized protocol for I_Ks_ with Kernik-Clancy current response.

**Appendix – Figure 8.**
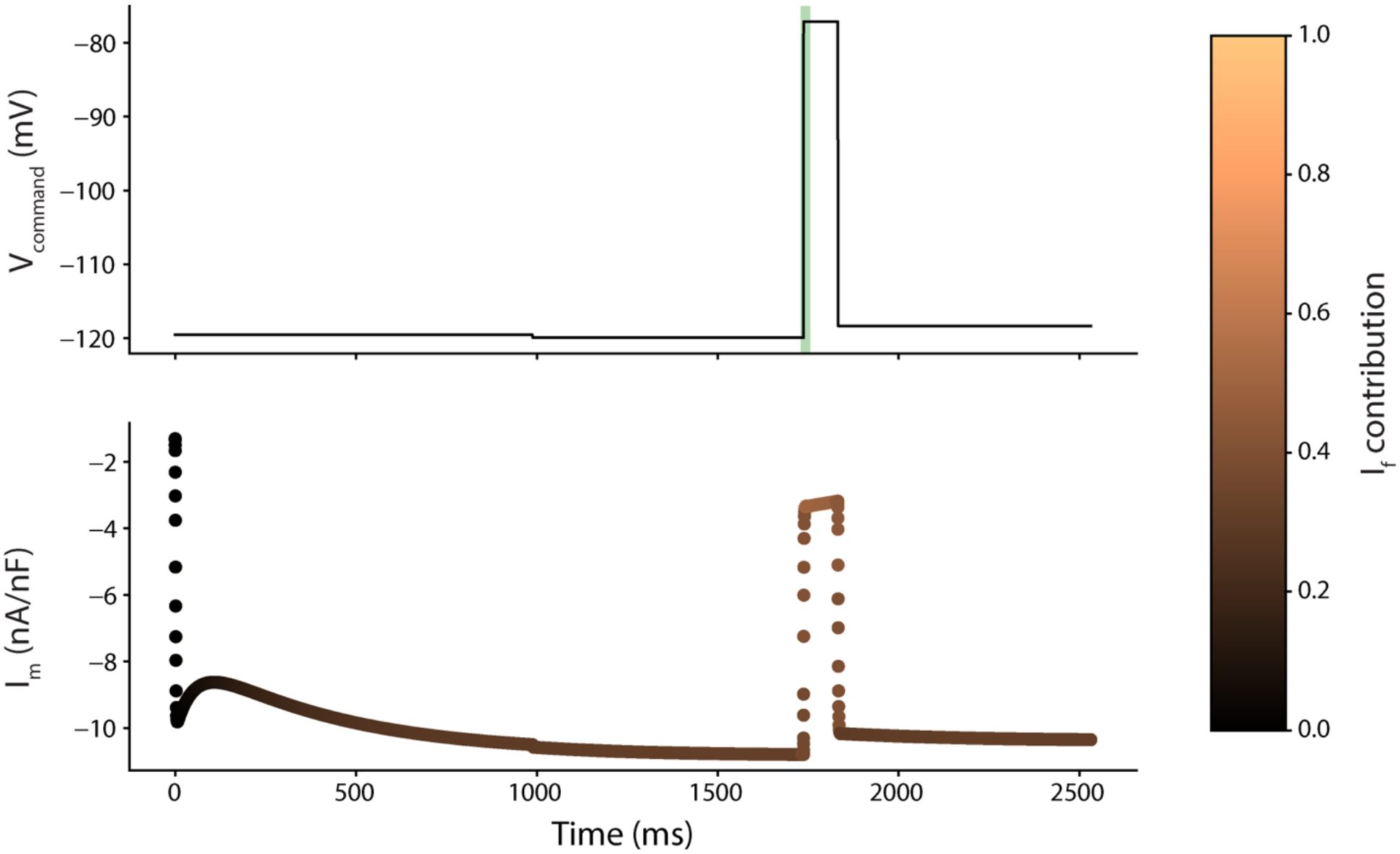
The optimized protocol for I_f_ with Kernik-Clancy current response.

**Appendix – Figure 9.**
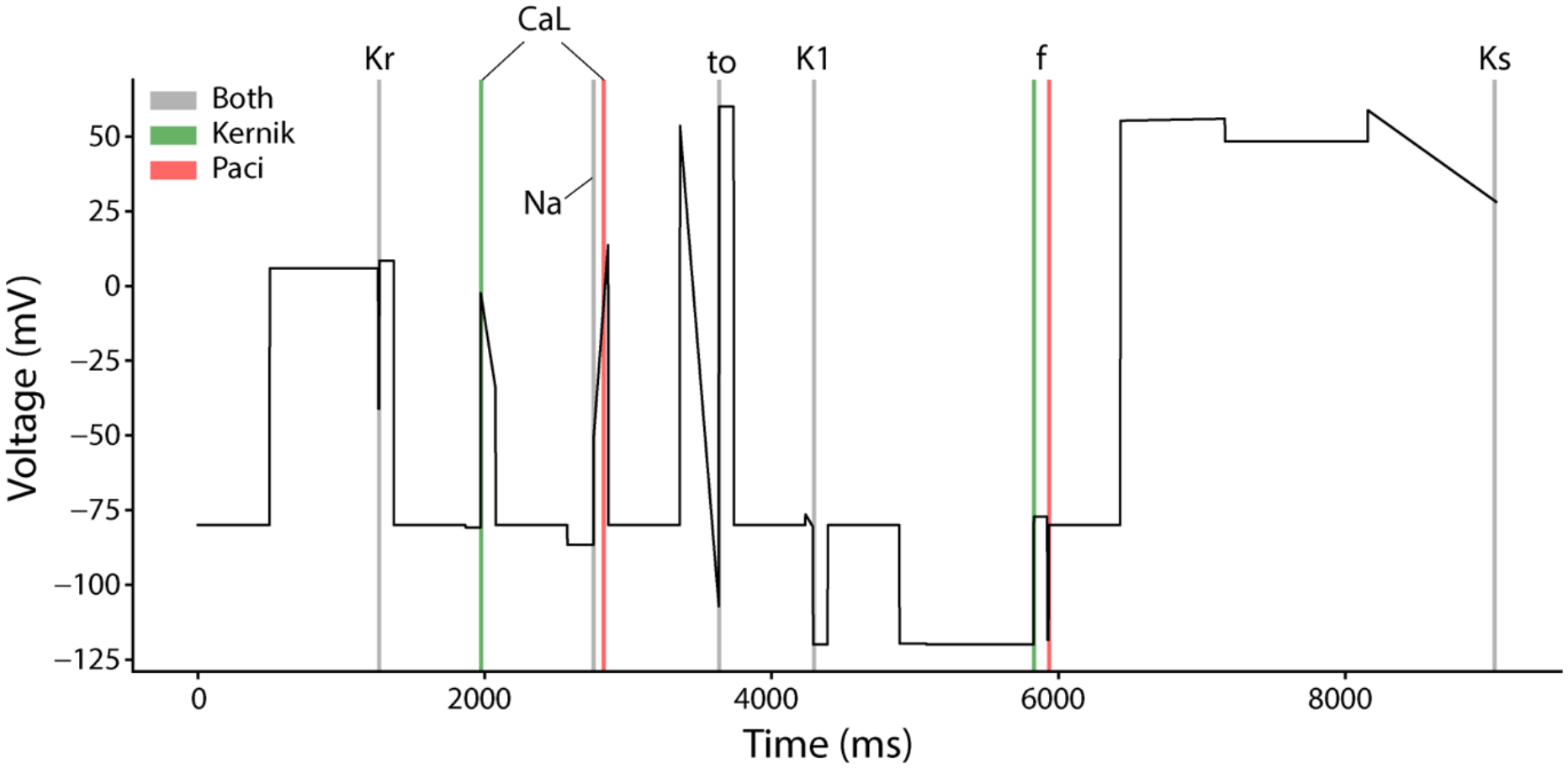
The timepoints of maximum current isolation for the Paci and Kernik-Clancy models. Five (I_K1_, I_to_, I_Kr_, I_Ks_, and I_Na_) of the seven currents in the Paci model were isolated within 10 ms of when they were isolated in the Kernik-Clancy model. These time windows are highlighted grey in the figure. The maximum I_CaL_ isolation in the Paci model occurs far from where the current is maximized in the Kernik model. However, the Paci model had a current isolation within 5% of its maximum during the Kernik-Clancy window. The timepoints for I_f_ also differed between the two models. However, these timepoints are near one another and have similar voltage dynamics, indicating that the Kernik-Clancy timepoint is likely generalizable for these currents.

**Appendix – Figure 10.**
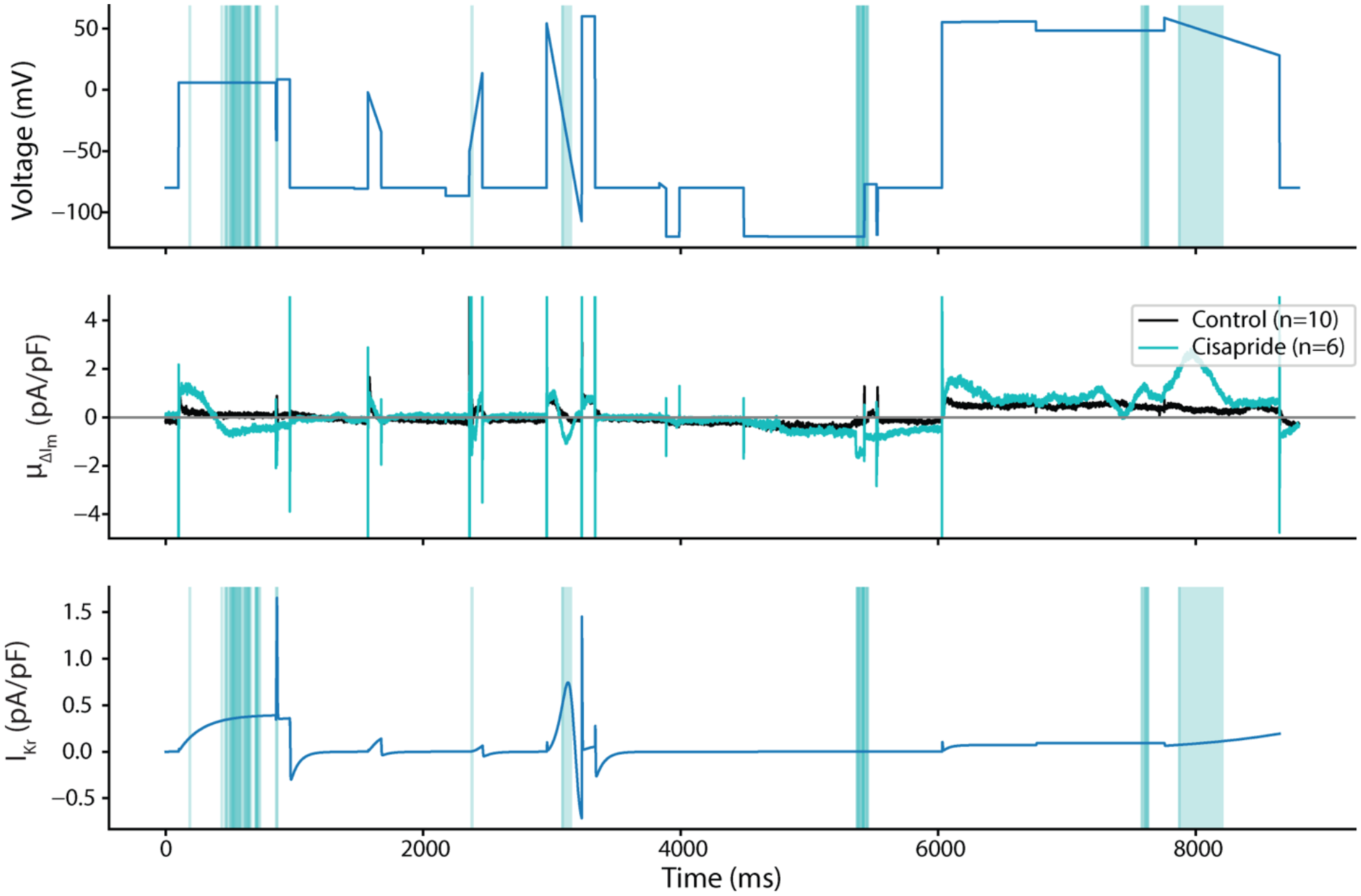
Differences in cell response to cisapride vs. DMSO. The voltage clamp protocol (top), average change in drug response from pre- to post-drug application for both DMSO and cisapride (middle), and the Kernik-Clancy I_Kr_ response to the voltage clamp protocol (bottom). The blue overlays indicate where there is a significant difference (p<.05) between the average cisapride and DMSO responses. We expected cisapride to strongly and specifically block I_Kr_. The bottom panel shows that most of the areas that are significantly different occur when I_Kr_ is present in the Kernik-Clancy model.

**Appendix – Figure 11.**
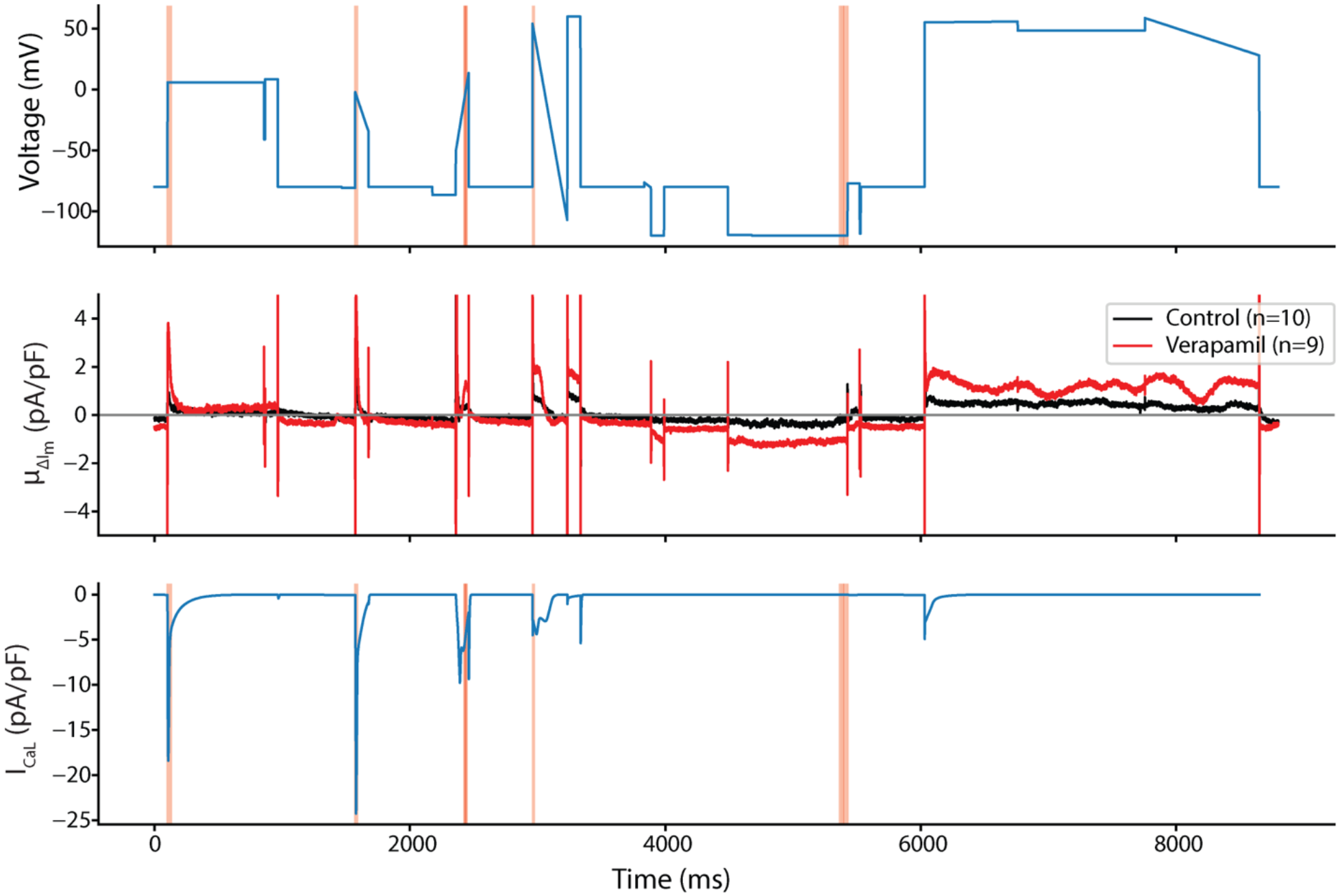
Differences in cell response to verapamil vs. DMSO. This figure shows the voltage clamp protocol (top), average change in drug response from pre- to post-drug application for both DMSO and verapamil (middle), and the Kernik-Clancy I_CaL_ response to the voltage clamp protocol. The red overlays indicate where there is a significant difference (p<.05) between the average verapamil and DMSO responses. At the concentration tested, we expect verapamil to block ∼40% of I_CaL_ and ∼20% of I_Kr_. The bottom panel shows that most of the areas that are significantly different occur when the I_CaL_ is present in the Kernik-Clancy model. There are two brief windows that the functional t-test identifies after 4000 ms, that are not likely I_Kr_ or I_CaL_.

**Appendix – Figure 12.**
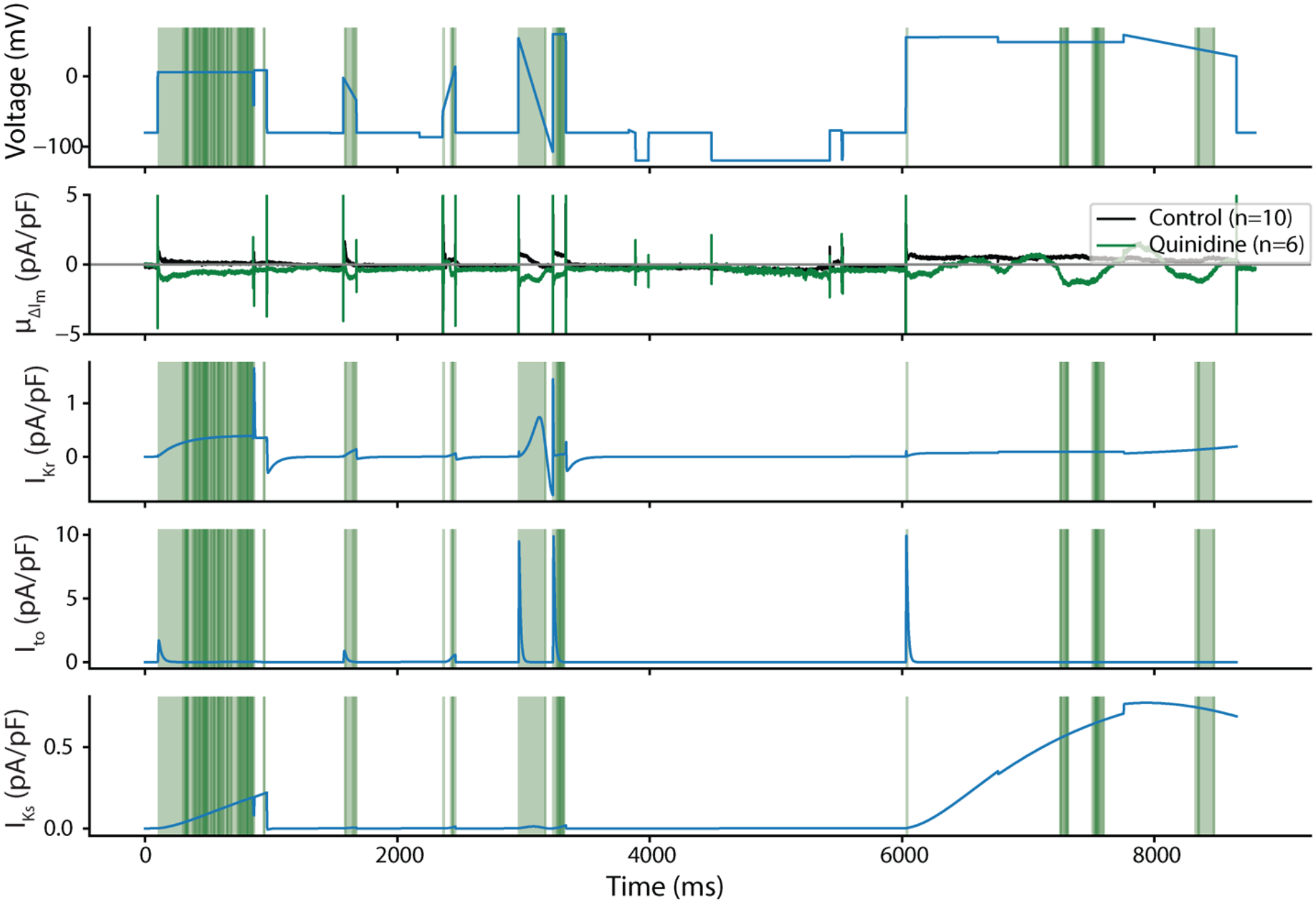
Differences in cell response to quinidine vs. DMSO. This figure shows the voltage clamp protocol, average change in drug response from pre- to post-drug application for both DMSO and quinidine, and the Kernik-Clancy I_Kr_, I_to_, and I_Ks_ responses to the voltage clamp protocol. The green overlays indicate where there is a significant difference (p<.05) between the average quinidine and DMSO responses. At the concentration tested, we expect quinidine to block ∼89% of I_Kr_, ∼43% of I_to_, and ∼27% of I_Ks_. The significance windows overlap very well with the Kernik-Clancy I_Kr_, I_to_, and I_Ks_ currents. This is to be expected, as quinidine is known to be a strong and general blocker of potassium currents.

**Appendix – Figure 13.**
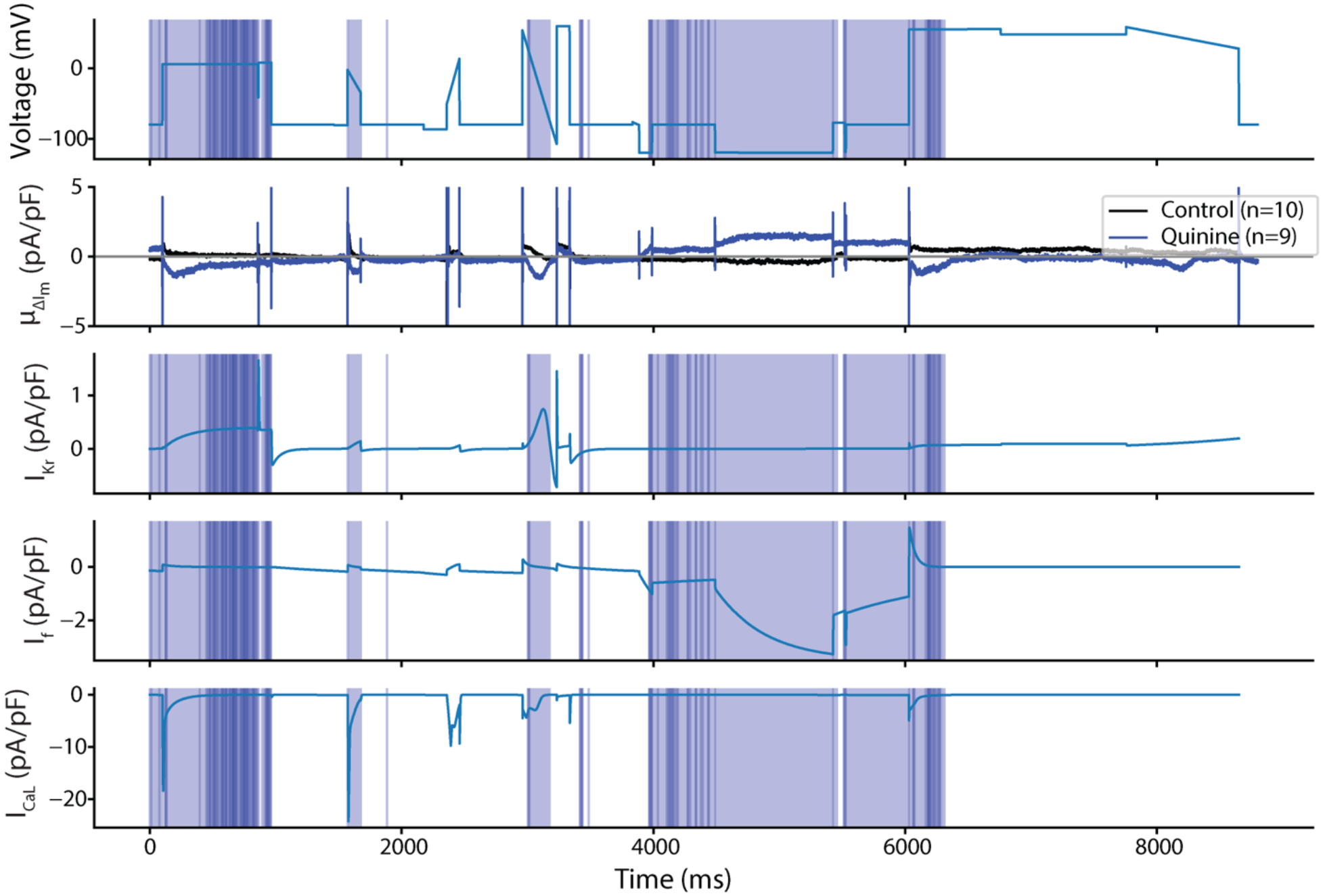
Differences in cell response to quinine vs. DMSO. This figure shows the voltage clamp protocol, average change in drug response from pre- to post-drug application for both DMSO and quinidine, and the Kernik-Clancy I_Kr_, I_f_, and I_CaL_ responses to the voltage clamp protocol. The blue overlays indicate where there is a significant difference (p<.05) between the average quinine and DMSO responses. At the concentration tested, we expect quinine to block ∼72% of I_Kr_ and ∼29% of I_CaL_. During the experiments, we noticed a likely block of I_f_ with quinine treatment. In figure 7, we show how we calculate a block of ∼32% of I_f_ by quinine at this concentration using a HEK-HCN1 cell line. The significance windows overlap very well with the Kernik-Clancy I_Kr_, I_CaL_, and I_f_ currents.

**Appendix – Figure 14.**
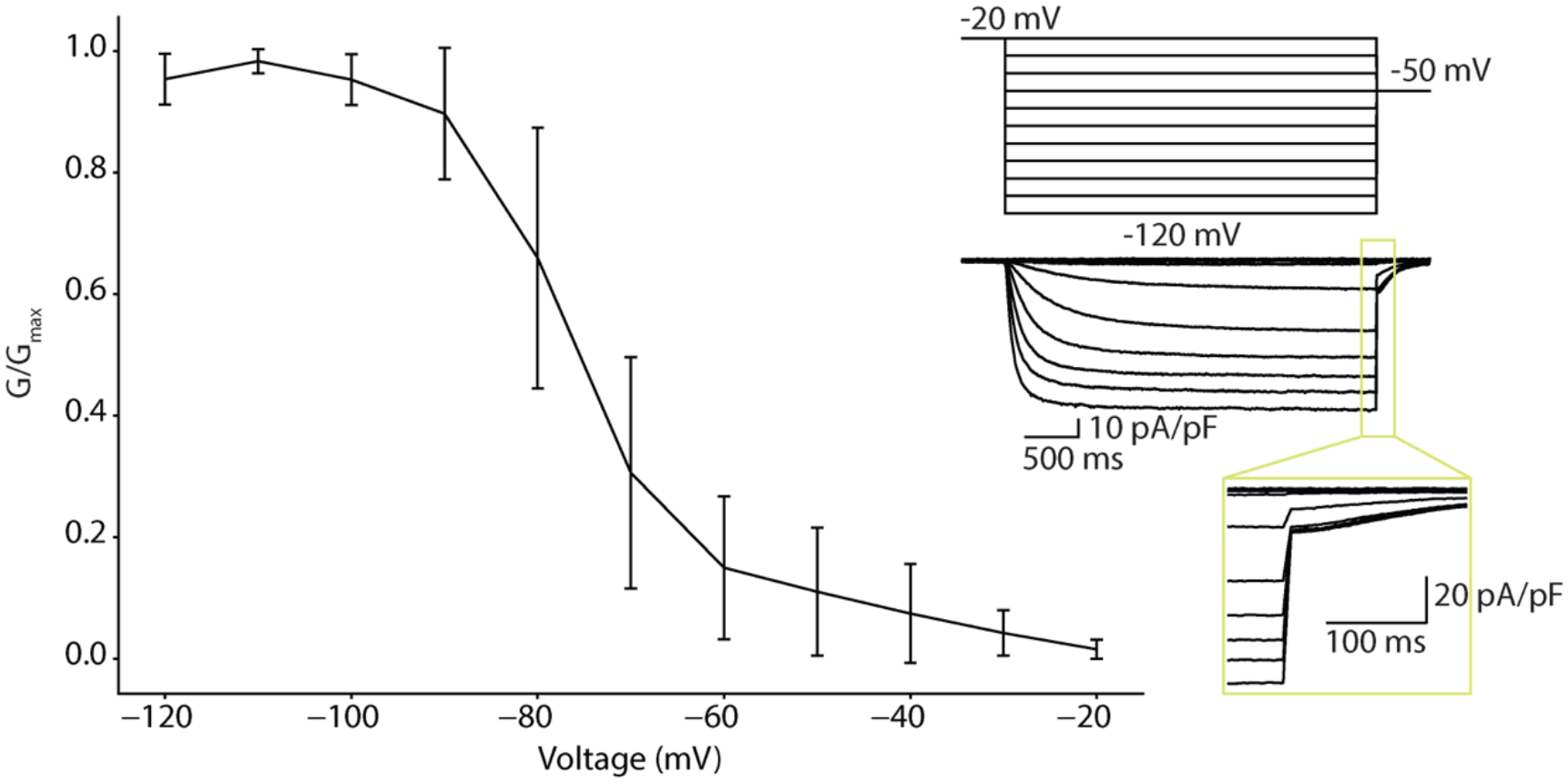
Max current vs voltage for HCN1 tail current. The max conductance-voltage curve was found by stepping to - 50mV after the channels had been activated with a depolarizing step. The max tail current values in this plot indicate that most, if not all, funny current channels are open when stepping to voltages below −100mV.

## Notes

### Competing Interest Statement

The authors have declared no competing interest.

## REFERENCES

Beattie KA, Hill AP, Bardenet R, Cui Y, Vandenberg JI, Gavaghan DJ, de Boer TP, Mirams GR. 2018. Sinusoidal voltage protocols for rapid characterisation of ion channel kinetics. J Physiol 596:1813–1828. doi:10.1113/JP275733

Bedut S, Seminatore-Nole C, Lamamy V, Caignard S, Boutin JA, Nosjean O, Stephan JP, Coge F. 2016. High-throughput drug profiling with voltage-and calcium-sensitive fluorescent probes in human iPSC-derived cardiomyocytes. Am J Physiol - Hear Circ Physiol 311:H44– H53. doi:10.1152/ajpheart.00793.2015

Britton OJ, Abi-Gerges N, Page G, Ghetti A, Miller PE, Rodriguez B. 2017. Quantitative comparison of effects of dofetilide, sotalol, quinidine, and verapamil between human ex vivo trabeculae and in silico ventricular models incorporating inter-individual action potential variability. Front Physiol 8:1–19. doi:10.3389/fphys.2017.00597

Burridge PW, Matsa E, Shukla P, Lin ZC, Churko JM, Ebert AD, Lan F, Diecke S, Huber B, Mordwinkin NM, Plews JR, Abilez OJ, Cui B, Gold JD, Wu JC, Paul Burridge or W. 2014. Chemically Defined and Small Molecule-Based Generation of Human Cardiomyocytes HHS Public Access. Nat Methods 11:855–860. doi:10.1038/nmeth.2999.Chemically

Charrez B, Charwat V, Siemons B, Finsberg H, Miller EW, Edwards AG, Healy KE. 2021. In vitro safety “clinical trial” of the cardiac liability of drug polytherapy. Clin Transl Sci 14:1155– 1165. doi:10.1111/cts.13038

Churko JM, Burridge PW, Wu JC. 2013. Cellular Cardiomyoplasty: Methods and Protocols, Methods in Molecular Biology 1036:81–88. doi:10.1007/978-1-62703-511-8

Costabal FS, Matsuno K, Yao J, Perdikaris P, Kuhl E. 2019. Machine learning in drug development: Characterizing the effect of 30 drugs on the QT interval using Gaussian process regression, sensitivity analysis, and uncertainty quantification. Comput Methods Appl Mech Eng 348:313–333. doi:10.1016/j.cma.2019.01.033

Crumb WJ, Vicente J, Johannesen L, Strauss DG. 2016. An evaluation of 30 clinical drugs against the comprehensive in vitro proarrhythmia assay (CiPA) proposed ion channel panel. J Pharmacol Toxicol Methods 81:251–262. doi:10.1016/j.vascn.2016.03.009

De Bruin ML, Pettersson M, Meyboom RHB, Hoes AW, Leufkens HGM. 2005. Anti-HERG activity and the risk of drug-induced arrhythmias and sudden death. Eur Heart J 26:590–597. doi:10.1093/eurheartj/ehi092

Demarco KR, Yang P-C, Singh V, Furutani K, Dawson J, Jeng M-T, Fettinger J, Bekker S, Ngo V, Noskov S, Yarov-Yarovoy V, Sack J, Wulff H, Clancy C, Vorobyov I. 2020. Molecular determinants of pro-arrhythmia proclivity of d- and l-sotalol via a multi-scale modeling pipeline. J Mol Cell Cardiol 115800. doi:10.1016/j.yjmcc.2021.05.015

EMA. 2005. ICH S7B Note for Guidance on the Nonclinical Evaluation of the Potential for Delayed Ventricular Repolarization (QT Interval Prolongation) by Human Pharmaceuticals. Int Conf Harmon Tech Requir Regist Pharm Hum Use.

Fabbri A, Goversen B, Vos MA, van Veen TAB, de Boer TP. 2019. Required GK1 to Suppress Automaticity of iPSC-CMs Depends Strongly on IK1 Model Structure. Biophys J 117:2303– 2315. doi:10.1016/j.bpj.2019.08.040

Fortin FA, De Rainville FM, Gardner MA, Parizeau M, Gagńe C. 2012. DEAP: Evolutionary algorithms made easy. J Mach Learn Res 13:2171–2175.

Garg P, Oikonomopoulos A, Chen H, Li Y, Lam CK, Sallam K, Perez M, Lux RL, Sanguinetti MC, C JCW. 2019. Genome Editing and Induced Pluripotent Stem Cells in Cardiac Channelopathy. J Am Coll Cardiol 72:62–75. doi:10.1016/j.jacc.2018.04.041.Genome

Giannetti F, Benzoni P, Campostrini G, Milanesi R, Bucchi A. 2021. A detailed characterization of the hyperpolarization - activated “funny” current (If) in human-induced pluripotent stem cell (iPSC)–derived cardiomyocytes with pacemaker activity. Pflügers Arch - Eur J Physiol. doi:10.1007/s00424-021-02571-w

Gintant G. 2011. An evaluation of hERG current assay performance: Translating preclinical safety studies to clinical QT prolongation. Pharmacol Ther 129:109–119. doi:10.1016/j.pharmthera.2010.08.008

Gong JQX, Sobie EA. 2018. Population-based mechanistic modeling allows for quantitative predictions of drug responses across cell types. npj Syst Biol Appl 4. doi:10.1038/s41540-018-0047-2

Goversen B, Becker N, Stoelzle-Feix S, Obergrussberger A, Vos MA, van Veen TAB, Fertig N, de Boer TP. 2018a. A hybrid model for safety pharmacology on an automated patch clamp platform: Using dynamic clamp to join iPSC-derived cardiomyocytes and simulations of Ik1 ion channels in real-time. Front Physiol 8:1–10. doi:10.3389/fphys.2017.01094

Goversen B, van der Heyden MAG, van Veen TAB, de Boer TP. 2018b. The immature electrophysiological phenotype of iPSC-CMs still hampers in vitro drug screening: Special focus on IK1. Pharmacol Ther 183:127–136. doi:10.1016/j.pharmthera.2017.10.001

Groenendaal W, Ortega FA, Kherlopian AR, Zygmunt AC, Krogh-Madsen T, Christini DJ. 2015. Cell-Specific Cardiac Electrophysiology Models. PLoS Comput Biol 11:1–22. doi:10.1371/journal.pcbi.1004242

Hancox JC, McPate MJ, El Harchi A, Zhang Y hong. 2008. The hERG potassium channel and hERG screening for drug-induced torsades de pointes. Pharmacol Ther 119:118–132. doi:10.1016/j.pharmthera.2008.05.009

Hobbs KH, Hooper SL. 2008. Using complicated, wide dynamic range driving to develop models of single neurons in single recording sessions. J Neurophysiol 99:1871–1883. doi:10.1152/jn.00032.2008

Hodgkin AL, Huxley AF. 1952. A quantitative description of membrane current and its application to conduction and excitation in nerve. J Physiol 117:500–44. doi:10.1113/jphysiol.1952.sp004764

Ishihara K, Sarai N, Asakura K, Noma A, Matsuoka S. 2009. Role of Mg2+ block of the inward rectifier K+ current in cardiac repolarization reserve: A quantitative simulation. J Mol Cell Cardiol 47:76–84. doi:10.1016/j.yjmcc.2009.03.008

Jæger KH, Charwat V, Wall S, Healy KE, Tveito A. 2021a. Identifying Drug Response by Combining Measurements of the Membrane Potential, the Cytosolic Calcium Concentration, and the Extracellular Potential in Microphysiological Systems. Front Pharmacol 11:1–16. doi:10.3389/fphar.2020.569489

Jæger KH, Wall S, Tveito A. 2021b. Computational prediction of drug response in short QT syndrome type 1 based on measurements of compound effect in stem cell-derived cardiomyocytes. PLoS Comput Biol 17:e1008089. doi:10.1371/journal.pcbi.1008089

Johannesen L, Vicente J, Mason JW, Sanabria C, Waite-Labott K, Hong M, Guo P, Lin J, Sørensen JS, Galeotti L, Florian J, Ugander M, Stockbridge N, Strauss DG. 2014. Differentiating drug-induced multichannel block on the electrocardiogram: Randomized study of dofetilide, quinidine, ranolazine, and verapamil. Clin Pharmacol Ther 96:549–558. doi:10.1038/clpt.2014.155

Kargol A. 2013. Wavelet-based protocols for ion channel electrophysiology. BMC Biophys 6. doi:10.1186/2046-1682-6-3

Kargol A, Hosein-Sooklal A, Constantin L, Przestalski M. 2004. Application of oscillating potentials to the Shaker potassium channel. Gen Physiol Biophys 23:53–75.

Kernik DC, Morotti S, Wu H Di, Garg P, Duff HJ, Kurokawa J, Jalife J, Wu JC, Grandi E, Clancy CE. 2019. A computational model of induced pluripotent stem-cell derived cardiomyocytes incorporating experimental variability from multiple data sources. J Physiol 597:4533– 4564. doi:10.1113/JP277724

Kernik DC, Yang PC, Kurokawa J, Wu JC, Clancy CE. 2020. A computational model of induced pluripotent stem-cell derived cardiomyocytes for high throughput risk stratification of KCNQ1 genetic variants. PLoS Comput Biol 16:1–28. doi:10.1371/JOURNAL.PCBI.1008109

Keser IK. 2014. Comparing two mean humidity curves using functional t-tests: Turkey case. Electron J Appl Stat Anal 7:254–278. doi:10.1285/i20705948v7n2p254

Klimas A, Ambrosi CM, Yu J, Williams JC, Bien H, Entcheva E. 2016. OptoDyCE as an automated system for high-throughput all-optical dynamic cardiac electrophysiology. Nat Commun 7:1–12. doi:10.1038/ncomms11542

Kopljar I, Lu HR, Van Ammel K, Otava M, Tekle F, Teisman A, Gallacher DJ. 2018. Development of a Human iPSC Cardiomyocyte-Based Scoring System for Cardiac Hazard Identification in Early Drug Safety De-risking. Stem Cell Reports 11:1365–1377. doi:10.1016/j.stemcr.2018.11.007

Lasser KE, Allen PD, Woolhandler SJ, Himmelstein DU, Wolfe SM, Bor DH. 2002. Timing of new black box warnings and withdrawals for prescription medications. J Am Med Assoc 287:2215–2220. doi:10.1001/jama.287.17.2215

Lei CL, Clerx M, Whittaker DG, Gavaghan DJ, de Boer TP, Mirams GR. 2020. Accounting for variability in ion current recordings using a mathematical model of artefacts in voltage-clamp experiments. Philos Trans A Math Phys Eng Sci 378:20190348. doi:10.1098/rsta.2019.0348

Li W, Luo X, Ulbricht Y, Wagner M, Piorkowski C, El-Armouche A, Guan K. 2019. Establishment of an automated patch-clamp platform for electrophysiological and pharmacological evaluation of hiPSC-CMs. Stem Cell Res 41:101662. doi:10.1016/j.scr.2019.101662

Lu HR, Zeng H, Kettenhofen R, Guo L, Kopljar I, van Ammel K, Tekle F, Teisman A, Zhai J, Clouse H, Pierson J, Furniss M, Lagrutta A, Sannajust F, Gallacher DJ. 2019. Assessing Drug-Induced Long QT and Proarrhythmic Risk Using Human Stem-Cell-Derived Cardiomyocytes in a Ca2+ Imaging Assay: Evaluation of 28 CiPA Compounds at Three Test Sites. Toxicol Sci 170:345–356. doi:10.1093/toxsci/kfz102

Mathur A, Loskill P, Shao K, Huebsch N, Hong SG, Marcus SG, Marks N, Mandegar M, Conklin BR, Lee LP, Healy KE. 2015. Human iPSC-based cardiac microphysiological system for drug screening applications. Sci Rep 5:1–7. doi:10.1038/srep08883

Millonas MM, Hanck DA. 1998. Nonequilibrium response spectroscopy of voltage-sensitive ion channel gating. Biophys J 74:210–229. doi:10.1016/S0006-3495(98)77781-1

Paci M, Pölönen RP, Cori D, Penttinen K, Aalto-Setälä K, Severi S, Hyttinen J. 2018. Automatic optimization of an in silico model of human iPSC derived cardiomyocytes recapitulating calcium handling abnormalities. Front Physiol 9:1–14. doi:10.3389/fphys.2018.00709

Passini E, Britton OJ, Lu HR, Rohrbacher J, Hermans AN, Gallacher DJ, Greig RJH, Bueno-Orovio A, Rodriguez B. 2017. Human in silico drug trials demonstrate higher accuracy than animal models in predicting clinical pro-arrhythmic cardiotoxicity. Front Physiol 8:1–15. doi:10.3389/fphys.2017.00668

Pioner JM, Santini L, Palandri C, Martella D, Lupi F, Langione M, Querceto S, Grandinetti B, Balducci V, Benzoni P, Landi S, Barbuti A, Lupi FF, Boarino L, Sartiani L, Tesi C, Mack DL, Regnier M, Cerbai E, Parmeggiani C, Poggesi C, Ferrantini C, Coppini R. 2019. Optical investigation of action potential and calcium handling maturation of hiPSC-cardiomyocytes on biomimetic substrates. Int J Mol Sci 20. doi:10.3390/ijms20153799

Quach B, Krogh-Madsen T, Entcheva E, Christini DJ. 2018. Light-Activated Dynamic Clamp Using iPSC-Derived Cardiomyocytes. Biophys J 115:2206–2217. doi:10.1016/j.bpj.2018.10.018

Roden DM. 2005. Drug-Induced Prolongation of the QT Interval. N Engl J Med 350:1013. doi:10.1097/01.sa.0000158587.83528.53

Sager PT, Gintant G, Turner JR, Pettit S, Stockbridge N. 2014. Rechanneling the cardiac proarrhythmia safety paradigm: A meeting report from the Cardiac Safety Research Consortium. Am Heart J 167:292–300. doi:10.1016/j.ahj.2013.11.004

Sahli-Costabal F, Seo K, Ashley E, Kuhl E. 2020. Classifying Drugs by their Arrhythmogenic Risk Using Machine Learning. Biophys J 118:1165–1176. doi:10.1016/j.bpj.2020.01.012

Sutanto H, Heijman J. 2020. Beta-Adrenergic Receptor Stimulation Modulates the Cellular Proarrhythmic Effects of Chloroquine and Azithromycin. Front Physiol 11. doi:10.3389/fphys.2020.587709

Tomek J, Bueno-Orovio A, Passini E, Zhou X, Minchole A, Britton O, Bartolucci C, Severi S, Shrier A, Virag L, Varro A, Rodriguez B. 2019. Development, calibration, and validation of a novel human ventricular myocyte model in health, disease, and drug block. Elife 8:1–47. doi:10.7554/eLife.48890

Tveito A, Jæger KH, Huebsch N, Charrez B, Edwards AG, Wall S, Healy KE. 2018. Inversion and computational maturation of drug response using human stem cell derived cardiomyocytes in microphysiological systems. Sci Rep 8:1–14. doi:10.1038/s41598-018-35858-7

Varshneya M, Irurzun-Arana I, Campana C, Dariolli R, Gutierrez A, Pullinger TK, Sobie EA. 2021. Investigational Treatments for COVID-19 may Increase Ventricular Arrhythmia Risk Through Drug Interactions. CPT pharmacometrics Syst Pharmacol 10:100–107. doi:10.1002/psp4.12573

Whittaker DG, Capel RA, Hendrix M, Chan XHS, Herring N, White NJ, Mirams GR, Burton R-AB. 2021. Cardiac TdP risk stratification modelling of anti-infective compounds including chloroquine and hydroxychloroquine. R Soc Open Sci 8. doi:10.1098/rsos.210235

Windley MJ, Lee W, Vandenberg JI, Hill AP. 2018. The Temperature Dependence of Kinetics Associated with Drug Block of hERG Channels Is Compound-Specific and an Important Factor for Proarrhythmic Risk Prediction. Mol Pharmacol 94:760–769. doi:10.1124/mol.117.111534

Yang PC, Demarco KR, Aghasafari P, Jeng MT, Dawson JRD, Bekker S, Noskov SY, Yarov-Yarovoy V, Vorobyov I, Clancy CE. 2020. A Computational Pipeline to Predict Cardiotoxicity: From the Atom to the Rhythm. Circ Res 947–964. doi:10.1161/CIRCRESAHA.119.316404

Zhou X, Qu Y, Passini E, Bueno-Orovio A, Liu Y, Vargas HM, Rodriguez B. 2020. Blinded in silico drug trial reveals the minimum set of ion channels for torsades de pointes risk assessment. Front Pharmacol 10:1–17. doi:10.3389/fphar.2019.01643

Zou L, Xue Y, Jones M, Heinbockel T, Ying M, Zhan X. 2018. The Effects of Quinine on Neurophysiological Properties of Dopaminergic Neurons. Neurotox Res 34:62–73. doi:10.1007/s12640-017-9855-1

